# Regulatory Feedbacks on Receptor and Non-receptor Synthesis for Robust Signaling

**DOI:** 10.1101/791509

**Authors:** A.D. Lander, Q. Nie, C. Sanchez-Tapia, A. Simonyan, F.Y.M. Wan

## Abstract

Elaborate feedback regulatory processes are thought to make biological developments robust, i.e., resistant to changes induced by genetic or environmental perturbations. How this might be done is still not completely understood. Previous numerical simulations on reaction-diffusion models of Dpp gradients in Drosophila wing imaginal disc showed that feedback (of the Hill’s function type) on (signaling) receptors and/or non-(signaling) receptors are of limited effectiveness in promoting robustness. Spatial nonuniformity of the feedback processes is thought to lead to serious shape distortion and a principal cause for ineffectiveness. Through mathematical modeling of a spatially uniform nonlocal feedback mechanism, the present paper provides a theoretical support of these observations. More significantly, the new approach also enables us to uncover in this paper a new, theory-based multi-feedback instrument for broadly effective promotion of robust signaling gradients.

## 1 Introduction

At some (early) stage of embryonic development of a biological organism, one or more proteins (known as morphogens or ligands) responsible for cell differentiation are synthesized and transported away from their sources to be bound to relevant cell receptors at different locations to form signaling morphogen-receptor complexes, known as a *signaling (spatial) gradients*. Such signaling gradients convey positional information for cells to adopt differential fates to result in tissue patterns. This process of cell differentiation is well established in developmental biology. For example, the morphogen *Decapentaplegic* (Dpp) involved in the development of the Drosophila wing imaginal disc is synthesized in a narrow region of about two-cell width at the boundary between the anterior and posterior compartments of the disc. Dpp molecules produced are transported away from the localized source and degrade upon reaching the edge of the disc. Some Dpp units bind reversibly with the cell-surface signaling receptor *Thickvein* (Tkv) to form a spatial gradient of signaling morphogen concentration over the span of the wing imaginal disc. Graded differences in receptor occupancy at different locations underlie the signaling differences that ultimately lead cells down different paths of development [12, 15, 16, 28, 42]. Simple models of this process of gradient formation have been shown theoretically to produce a unique signaling gradient that is monotone and asymptotically stable with respect to small perturbations (see [23, 24] for example).

For normal biological development, it is important that signaling morphogen gradients not be easily altered by genetic or epigenetic (such as environmental) fluctuations that affect the constitution of the biological organism [33]. Experimental results (carried out by S. Zhou in A.D. Lander’s lab (see also [49]) show that Dpp synthesis rate doubles when the ambient temperature is increased by 5.9°*C*. With such an increase in Dpp synthesis rate, the simple models developed in [23, 24, 25] would predict an enhanced or (more commonly called) “ectopic” signaling gradient quantitatively and qualitatively different from that under the (lower) normal ambient temperature. Yet development of the wing imaginal disc generally does not change significantly with temperature changes of such magnitude. The insensitivity of system output to sustained alterations in input or system characteristics so necessary for normal development is often termed *robustness* of biological development. How this robustness requirement is met has been the subject of a number of recent studies [8]-[11],[18, 22, 29, 30, 34, 39, 40, 45].

Evidence exists that regulatory feedback processes play a role in rendering biological developments robust, i.e., resistant to changes induced by genetic or environmental perturbations [13, 36]. How this might be done is still not completely understood. Among the first attempts to determine mechanisms for attaining robust developments, a negative feedback on receptor synthesis rate was investigated in [26]. A Hill function type negative feedback was incorporated into the basic morphogen gradient model of [24] to reduce the synthesis rate of Tkv by an amount that depends on the ectopic signaling morphogen concentration. Some numerical simulations of the model show that robustness is not achieved for any of the 10^6^ combinations of system parameter values in a parameter space of 6-dimensions. A subsequent theoretical analysis delineated and confirmed theoretically the ineffectiveness of this negative feedback mechanism [19]. Briefly, a Hill type negative feedback reduces the receptor synthesis rate nonuniformly, disproportionately more so at locations of high signaling morphogen concentration. Such reduction generally leads to a modified gradient of different slope and convexity from the normal (wild-type) gradient. The theoretical results suggest that a spatially uniform negative feedback responding to some overall measure of ectopicity (such as the average impact of the local changes on the system) may be more effective [19]. This suggestion has led to the initiation of a new general approach to attain robustness by way of a feedback mechanism that is spatially uniform [21, 39, 40, 45].

With a view that most feedback mechanisms have the ultimate effect of reducing the morphogen available for binding with signaling receptors, a proof-of-concept prototype model for a spatially uniform negative feedback on morphogen synthesis rate was first investigated in [21]. The findings in that preliminary effort provided the impetus to investigate the efficacy of the same feedback instrument for other more realistic spatially uniform feedback mechanisms in [45]. In this paper, we refine the models on reducing the availability of morphogen concentration for binding investigated in [21, 45] by modeling explicitly one of the processes known to reduce the morphogen availability. Among the different ways to reduce Dpp concentration is their binding with other protein molecules to form morphogen complexes that do not signal for cell differentiation [4]. Such (non-signaling) companion proteins are known to exist for Dpp and other BMP family ligands. They include Notum [14], Nog (noggin) [31, 47, 50], Chd (chordin) [37, 48], Dad (daughter against dpp) [43], Dally (division abnormally delayed) [1, 20], FST (follistatin (FST) [3, 17, 35, 46], Sog (short gastrulation) [5, 32], and various heparan sulfate proteoglycans [6]. Collectively, they are called *non-receptors* since they bind with morphogens but the resulting bound morphogen complexes have no role in cell differentiation.

Effects of non-receptors was modeled and analyzed in [27] where we extend the simple wing disc morphogen model of [23, 24] to include a fixed concentration of cell-surface non-receptor (induced instantaneously at the onset of the genetic or epigenetic perturbations at time *t_e_*). This simplest model offered the first theoretical glimpse into the inhibiting effects of non-receptors on the formation and properties of steady state signaling morphogen-receptor gradients. Subsequently, large scale computational studies of non-receptors synthesized at a prescribed (perturbations-induced) fixedrateattime *t_e_* to absorb excessive Dpp concentration was carried out in [26]. Extensive numerical simulations spanning a 6-dimensional parameter space showed that less than 9% of gradients are of appropriate size and shape but with a mean *robustness index* (a numerical value to be defined in a later section as a measure of robustness of the ectopic gradient relative to the wild type) more than doubling that defined to be acceptably close to the wild-type gradient prior to the perturbations. Most gradients generated are not biologically realistic for cell differentiation (for reasons such as high receptor occupancy throughout the wing imaginal disc away from its distal edge). These observations have been validated theoretically in [19, 29, 44]. Adding negative feedback on receptor synthesis rate to *a modest concentration of non-receptors* was found in [26] to result in a slightly broader range of robust gradients that are biologically useful and robustness cannot be attained with higher non-receptor synthesis rates (than the receptor synthesis rate) with any level of negative feedback on receptor synthesis rate. Additional negative feedback on the prescribed non-receptor synthesis would only further degrade the ectopic gradient.

The theoretical results on a Hill function type feedback on receptor synthesis shows that such spatially nonuniform feedback distorts the shape of the ectopic gradient and suggest that a spatially uniform feedback mechanism may reduce such distortion. A robustness-index based feedback mechanism has been developed in [21, 45] for this purpose. While the improvements associated with the new feedback process applied to receptor synthesis rate (and a few other biological processes, such as the binding rate, and receptor-mediated degradation rate) was found insufficient for robust signaling, it enables us to uncover the benefits of appropriate multi-feedback mechanisms that are much more efficient in promoting robustness. In relation to the numerical simulations of [26], we report in this paper one specific application of the multi-feedback combination to show how ectopicity induced positive feedback on non-receptor synthesis may be combined with a similar negative feedback on receptor synthesis for a very effective instrument for promoting robustness in biologically realistic ranges of system parameter values. We do this by examining a set of models that are the counterparts of those investigated in [26] but now with our new spatially uniform feedback instrument instead of the Hill function type previously employed. One significant feature of our approach is that the new models admit explicit exact solutions for biologically realistic gradients so that our results are theoretically conclusive and do not rely on numerical simulations. How non-receptors may or may not promote robustness can be seen explicitly from the mathematical expressions in terms of known functions for the signaling gradient concentration of the different models.

## 2 The Simplest Extracellular Model of Dpp Gradient Formation

To understand better the results of numerical simulations of [26], we re-examine the same three approaches to robustness there but now by way of a spatially uniform feedback. For this purpose, we work with the normalized form of the one-dimensional extracellular model of the Dpp gradient formation of [24]. To simplify the analysis without sacrificing any of the essential characteristics of the biological processes involved, we may take the wild-type Dpp synthesis rate *V_L_*(*X, T*) to be uniform in the direction along the boundary *X* = −*X*_min_ between the anterior and posterior compartments of the wing imaginal disc. Here, *X* is distance in direction perpendicular to the between-compartment boundary with *X* spanning [−*X*_min_, *X*_max_], *X*_max_ being the edge of the posterior compartment. With Dpp synthesized at a uniform rate in a narrow strip (of a couple of cells width) −*X*_min_ < *X* < 0, we idealize the synthesis rate by

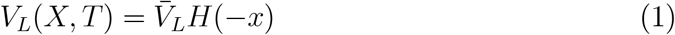

where *x* = *X*/*X*_max_. We also take the wild-type receptor synthesis rate *V_R_*(*X,T*) to be uniform throughout the posterior compartment with a steady state receptor concentration of

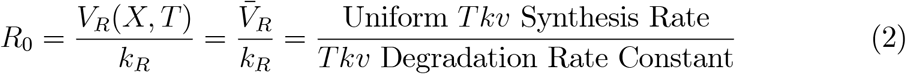

prior to the onset of Dpp synthesis at *T* = 0. Just as (the normalized)distance *x* in direction normal to the compartment boundary measured in units of the maximum distal width *X*_max_ of the posterior compartment, we also normalize the physical time *T* by setting 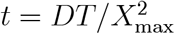 where *D* is the uniform diffusion coefficient.

Normal development of wing imaginal disc and other biological organisms may be altered by an enhanced morphogen synthesis rate stimulated by sustained genetic or epigenetic changes (in contrast to a one time perturbation of a existing steady state), starting at some time *t_e_* > 0. For example, *Dpp* synthesis rate in Drosophila imaginal disc has been shown to double when the ambient temperature is increased by 5.9°*C* (shown by S. Zhou while in A.D. Lander’s Lab). At a state of lower receptor occupancy, basic models for signaling gradient formation would have the corresponding steady state *ectopic* signaling ligand concentration increasing proportionately (see (19)-(20)) and its shape altered (and hence the cell fate at each spatial location as well) [24, 22]. Without the restriction of low receptor occupancy, these and other models have also shown the steady state ectopic signaling gradient to be an increasing function of synthesis rate, though not necessarily proportionately [24, 22, 19, 29].

Natural biological developments however are mostly unaffected by sustained environmental perturbations that should have altered them. To investigate possible mechanisms for managing possible ectopic developments induced by genetic or epigenetic perturbation, we introduce an ectopic (amplification) factor *e* ≥ 1 and work with a more general ligand synthesis rate 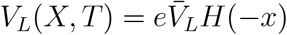 instead of (1). The basic extracellular model for Dpp gradient formation of [24] then consists of the following three normalized differential equations for the normalized concentrations of free Dpp concentration *a_e_*(*x, t*), Dpp-Tkv complexes concentration (or *signaling Dpp gradient* for short) *b_e_*(*x, t*), and the unoccupied Tkv *r_e_*(*x, t*), all measured in units of the steady state receptor concentration *R*_0_ introduced in (2):

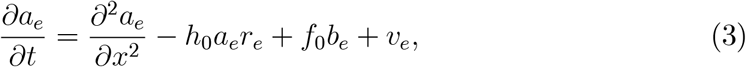

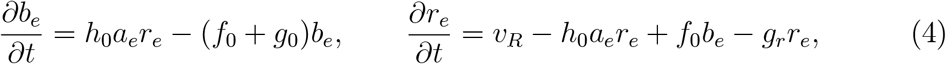

where the quantities *h*_0_, *g*_0_, and *f*_0_ are (per unit morphogen concentration) binding rate, receptor-mediated degradation rate and dissociation rate, all normalized by 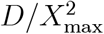. In the absence of feedback, the normalized Dpp and Tkv synthesis rates, *v_e_* and *v_R_*,are given by

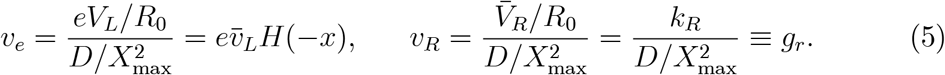

where *e* is the ligand synthesis amplification factor (or ectopicity) with *e* = 1 for the wild type.

The three differential equations are supplemented by the boundary conditions

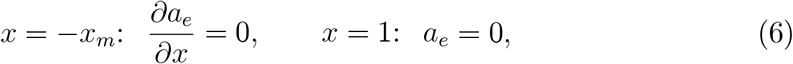

all for *t* > 0, and the initial conditions

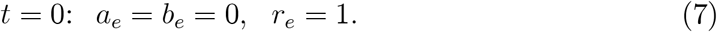

The *initial-boundary value problem* (IBVP) defined by (3)-(7) and its modified forms have been analyzed as mathematical models for ligand activities and tissue pattern formation in several of the references cited (e.g., [24, 19, 26]). Some basic results from [24] are summarized below for comparison with the new results on the effects of non-receptors to be analyzed herein.

### 2.1 Time Independent Steady State

Given that both the ligand and receptor synthesis rates are time independent, it has been shown in [24] that the extracellular model system (3)-(7) has a unique steady state,

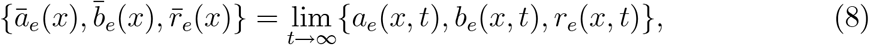

that is asymptotically stable with respect to small perturbations. With the three dependent variable not changing with time in steady state, the governing IBVP may be reduced to the following well-posed two-point boundary value problem (BVP) for *ā_e_*(*x*) [24]:

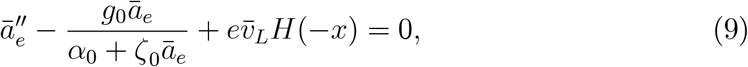

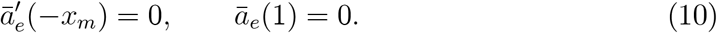

with

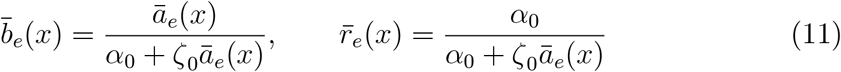

where

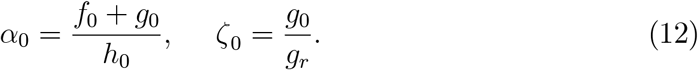

We note again that the ectopicity factor *e* is a constant for the level of abnormality in the ligand synthesis rate with *e* = 1 for the wild-type development [24].

For the signaling gradient to induce a distinct biological tissue pattern, it should *not* be nearly uniform over a significant spatial span of the solution domain (−*x_m_*, 1) as there would not be a pattern over that span. For this reason, the free morphogen concentration *ā_e_*(*x*) associated with a biologically realistic gradient system cannot be so large that

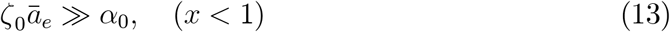

or (with *f*_0_ ≪ *g*_0_) *ā_e_* ≫ *g_r_*/*h*_0_. If the condition (13) should be met, we would have

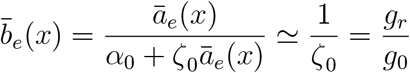

uniform except for a boundary layer adjacent to the edge *x* = 1. While *ζ*_0_*ā_e_*(*x*) = *O*(*α*_0_) is not the only requirement for a biologically realistic gradient system, we formulate it as a necessary condition for systems worthy of examination.

#### Criterion 1

*For a morphogen gradient system to induce a biological meaningful pattern, it free morphogen concentration must **not** be so large for ζ*_0_*ā_e_*(*x*) ≫ *α*_0_ *outside a narrow region adjacent to the edge x* = 1.

### 2.2 Low Receptor Occupancy

At the other extreme, when the free morphogen concentration *ā_e_*(*x*) is sufficiently low over the span of the solution interval (0,1) so that

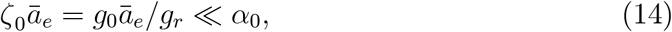

we may neglect terms involving *ζ*_0_*ā_e_* in (9)-(11) to get an approximate set of solutions {*A_e_*(*x*), *B_e_*(*x*), *R_e_*(*x*)} determined by the initial value problem (IVP)

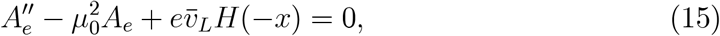

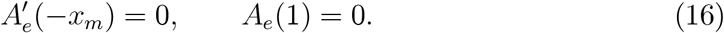

with

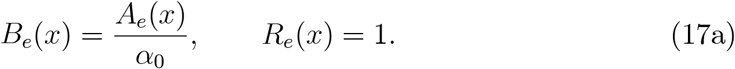

and

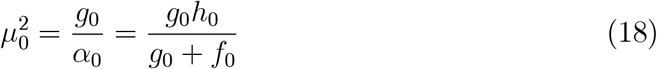

The exact solution for *A_e_*(*x*) = *eA*_1_(*x*) obtained previously in [24] is

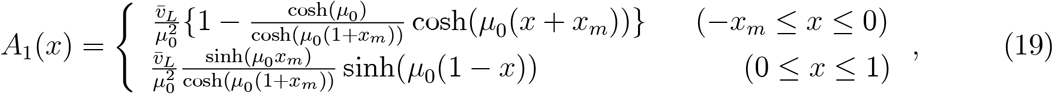

with

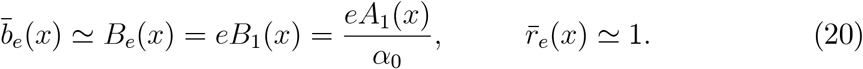

It would be natural to characterize a morphogen system with such low concentration of free morphogen so that (14) holds to be in a state of *low receptor occupancy* (LRO) since there are few free ligand available to bind with signaling receptors. However, if we have also *μ*_0_ ≫ 1, the expression for *A_e_*(*x*) in the signaling range of 0 ≤ *x* < 1 is effectively a boundary layer adjacent to the edge of the ligand production region, steep near *x* = 0 and dropping sharply to 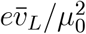 (which is rather small) away from that edge. Generally, the bound (signaling) morphogen gradient 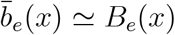 should change gradually if it is to lead to a distinct biological pattern. To limit our discussion to these biologically meaningful gradient systems, we adopt the following definition for a gradient system in a steady state of LRO:

#### Definition 2

*A morphogen system is in a steady state of **low receptor occupancy** (LRO) if the condition (14) is satisfied and μ*_0_ = *O*(1).

With the adoption of this definition, we may then restrict our attention mainly to LRO systems that give rise to distinctive biological tissue patterns. For such systems, the bound and free ligand concentrations change only gradually over their spatial span [0, *X*_max_].

For gradient systems for which neither (14) nor (13) is met, the following condition provides a criterion for identifying the kind of biological gradient systems worthy of investigation:

#### Criterion 3

*A gradient system is **not** biologically meaningful if μ*_0_ ≫ 1.

While Criterion 1 eliminates signaling gradients that are pretty much flat and with signaling receptors saturated away from the edge *x* = 1, Criterion 3 eliminates signaling gradients that are also pretty much flat but with signaling receptors sparsely occupied away from ligand synthesis region.

### 2.3 Root-Mean-Square Signaling Differential

We wish to make use of the extracellular model summarized in the preceding subsections to investigate the consequences of some feedback mechanisms on robust development to gain more insight into the results from numerical simulations obtained in [26]. We do this by working with a spatially uniform feedback to complement the conventional Hill function approach in modeling feedback processes. For this purpose, we need to have a quantitative robustness measure that quantifies succinctly deviation from the wild-type development. One such measure, the *root-mean-square signaling differential robustness index* (henceforth the *R_b_ robustness index* for short, or simply *robustness index* when there is no ambiguity) used in [22], is given below and applied to illustrate its effectiveness in measuring the ectopicity of a signaling gradient in a state of low receptor occupancy. This index is simpler to analyze compared to the *root-mean-square displacement differential robustness index R_x_* (previously denoted by *L* in [26]) which was also used earlier in [22] and will be examined in later sections of this paper.

The (*signal*) *robustness index R_b_* is the root mean square of the deviation of the ectopic signaling gradient *b_e_*(*x, t*) from wild-type signaling gradient *b*_1_(*x, t*):

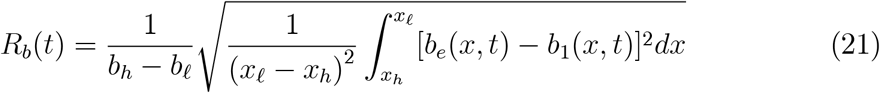

where 0 ≤ *b_ℓ_*(*t*) < *b_h_*(*t*) ≤ *b*_1_(−*x_m_, t*) and −*x_m_* ≤ *x_h_* < *x_ℓ_* ≤ 1. The quantities *x_ℓ_*, *x_h_*, *b_ℓ_* and *b_h_* may be chosen away from the extremities to minimize the exaggerated effects of outliers. For a system in steady state with

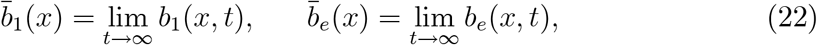

the robustness index *R_b_*(*t*) tends to a constant 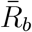:

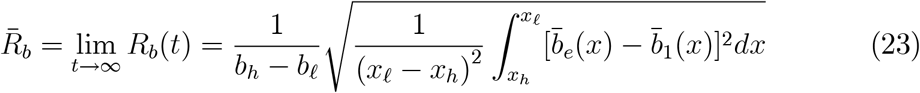

In subsequent developments, we set *x_h_* = 0 as the region of ligand synthesis [−*x_m_*, 0) is not expected to contribute significantly to signaling. We also take *x_ℓ_* = 1 so that *b_ℓ_*(1, *t*) = *b*_1_(1, *t*) = 0. For the case of low receptor occupancy, we take *b_h_* to be *B*_1_(0), the explicit LRO approximation of the steady state wild-type signaling gradient concentration value 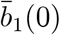, known from (19) and (20) to be

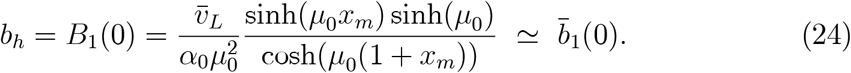

Having 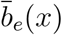 and 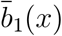 (by any numerical software for BVP of ODE), it is straightforward to evaluate the integral (23) to obtain 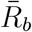 to see whether or not the ectopicity of the deformed gradient is still acceptable.

### 2.4 Approximate Robustness Index for LRO State

For a morphogen system in a state of LRO, we have from (19)-(20) the following approximate steady state solutions for the signaling gradients, *B_e_*(*x*), of the (environmentally or genetically) perturbed system:

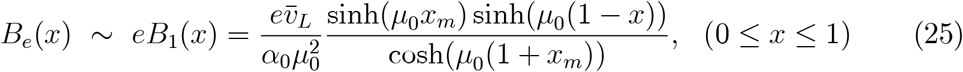

where 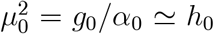 since *f*_0_ ≪ *g*_0_ for our model of the wing imaginal disc. The parameter *e* is the amplification factor of the ligand synthesis rate. Then the LRO approximation of 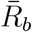 (with *x_ℓ_* = 1, *x_h_* = 0), denoted by *ρ*_0_, is given by

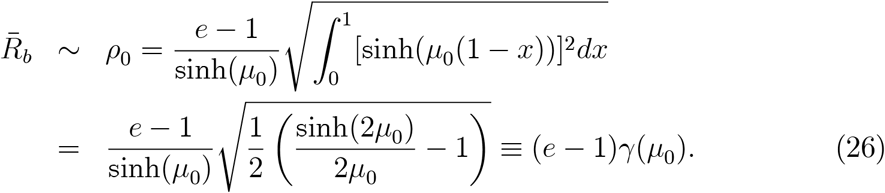

To be concrete and to make use of the finding of Zhou on the effect of a 5.9°*C* temperature change, we are mainly concerned with the *empirically observed case* of *e* = 2 in the subsequent development.

For a gradient system with *g*_0_ = 0.2, *f*_0_ = 0.001, *g_r_* = 1, *h*_0_ = 10, *x_m_* = 0.1 and 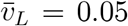 (corresponding to 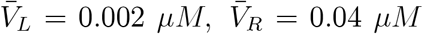, *D* = 10^-7^*cm*^2^/ sec., *X*_max_ = 0.01*cm*) with 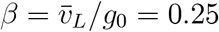 in Table 2 of [24], the steady state is in LRO state. For this case, the approximate solution for 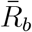 given by (26) is 0.3938… while the actual solution for 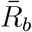 computed from an accurate numerical solution for the BVP for *ā*_2_(*x*) gives 0.3943… for a percentage error of less than 0.01%. If ligand synthesis rate is increased 20 times to 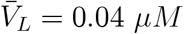, the percentage error of the low receptor occupancy approximation is still less than 1%. These comparisons serve to validate the numerical simulation code developed for exact numerical solutions of our model. (Consistent with the subsequent ease of analysis for the LRO and LRNO state, we have adopted the simplifying approximation *α*_0_ ≃ *g*_0_/*h*_0_ in the computation for both 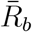 and *ρ*_0_ in this paper since *f*_0_ ≪ *g*_0_.)

**Table I.** Some Sample Solutions. *X*_max_ = 0.01 *cm*, *X*_min_ = 0.001 *cm*, *k_on_R*_0_ = 0.01/sec./μM, *k_deg_* = 2 × 10^−4^/sec., *k_R_* = 0.001/sec., *k_off_* = 10^−6^/sec., *e* = 2 *D* = 10^−7^ *cm*^2^/sec., 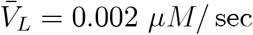., 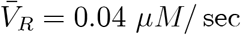.

|  |  |  |  |  |  |  |
| --- | --- | --- | --- | --- | --- | --- |
| $c$ | $\bar{R}_b$ | $\rho_c$ | $\bar{b}_e(0; c)$ | $eB_v(0)$ | $\bar{b}_1(0)$ | $B_1(0)$ |
| 0 | 0.3943 | 0.3938 | 0.1154 | 0.1169 | 0.0580 | 0.0584 |
| 0.5 | 0.3750 | 0.3736 | 0.1080 | 0.1096 | 0.0580 | 0.0584 |
| 1 | 0.3573 | 0.3565 | 0.1022 | 0.1062 | 0.0580 | 0.0584 |
| 2 | 0.3284 | 0.3284 | 0.0941 | 0.0959 | 0.0580 | 0.0584 |
| 4 | 0.2862 | 0.2872 | 0.0839 | 0.0857 | 0.0580 | 0.0584 |

**Table II.** (*g*_1_ = 0.2; *h*_1_ = 10, *f*_1_ = 0.01, *g_n_* = 10, *v_L_* = 0.05, *e* = 2)

| $(g_1 = 0.2; \quad h_1 = 10, \quad f_1 = 0.01, \quad g_n = 10; \quad v_L = 0.05, \quad e = 2)$ | | | | | | |
| --- | --- | --- | --- | --- | --- | --- |
| $Z$ | $\bar{R}_b$ | $\rho_Z$ | $\bar{b}_2(0, Z)$ | $eB_2(0; Z)$ | $\bar{b}_1(0, 0)$ | $B_1(0, 0)$ |
| 0 | 0.3943 | 0.3938 | 0.11517 | 0.1169 | 0.05790 | 0.0584 |
| 1 | 0.0714 | 0.0693 | 0.07398 | 0.0739 | 0.05790 | 0.0584 |
| 2 | 0.1298 | 0.1365 | 0.05597 | 0.0555 | 0.05790 | 0.0584 |

Our main interest however is in the use of the robustness index to induce an appropriate feedback mechanism for attaining robustness of signaling morphogen gradients. When an enhanced ligand system is not in a state of low receptor occupancy, the use of the approximate signaling robustness index based on the approximate solution (25) may not be sufficiently accurate. For these cases, numerical solutions of 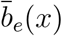 and the corresponding value for 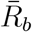 should be used instead of *B_e_*(*x*) and *ρ*_0_.

## 3 Feedback on Receptor Synthesis Rate

Excessive ligand concentration is known to down-regulate its own signaling receptor synthesis. In particular, Decapentaplegic (Dpp) represses the synthesis of its own receptor Tkv [28]. Another example is Wingless (Wg) repressing its signaling receptor DFz2 [7]. The down-regulation of Tkv by ectopic signaling Dpp gradient was modeled by a negative feedback of the Hill function type in [26] and was found to be ineffective as an instrument for robust development of the wing imaginal disc. A theoretical analysis of the model in [19] confirms the results of the numerical experiment and shows that the spatially nonuniform feedback distorts the shape of the output gradient as the feedback mechanism works to reduce its ectopicity. The observation suggests that a spatially uniform feedback mechanism may be more effective for ensuring robustness. We consider in [45] the originally (normalized) spatially uniform receptor synthesis rate 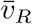 being down-regulated to *v_R_*(*t*) by a negative feedback factor *κ*^2^(*t*) that is a function of the signaling robustness index *R_b_*(*t*) in the form

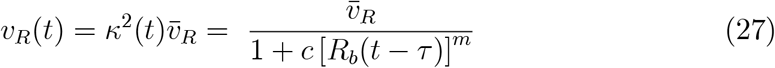

where the parameter *τ* corresponds to a possible time delay and {*c, m*} are two parameters to be chosen for appropriate feedback strength and sensitivity, respectively, similar to those for a Hill’s function.

It should be noted that the feedback process in (27) at any location does not depend on the ectopicity of the signaling gradient at that location, only on an average measure *R_b_*(*t*) of the excess over the span of the wing disc. Since it is not sensitive to the local environment of individual cell, it is less likely to contribute to the shape distortion of the resulting gradients. More detailed justification of such a non-local feedback mechanism can be found in [45].

For the investigation of the effects of our particular type of negative feedback on the receptor synthesis rate, we are interested in the modified signaling gradient (starting at *t* = 0) and the corresponding robustness index of the IBVP (3) - (7) but now with an enhanced ligand synthesis rate

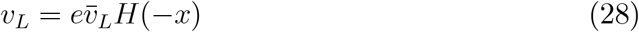

with an amplification factor *e* > 1 and a down-regulated receptor synthesis rate given by (27). In the presence of the feedback, the three ectopic gradients of the new IBVP are to be denoted by {*a_v_*(*x, t*), *b_v_*(*x, t*), *r_v_*(*x, t*)} ≡ {*a_e_*(*x, t*; *c*), *b_e_*(*x, t*; *c*), *r_v_*(*x, t*; *c*)} which reduce to {*a_e_*(*x, t*), *b_e_*(*x, t*), *r_e_*(*x, t*)} of the model without feedback (when *c* = 0). The corresponding robustness index is determined by

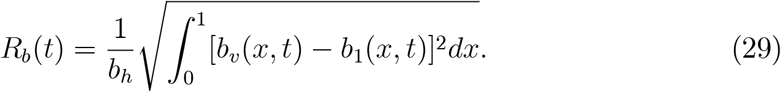

Unlike the situation in (21), *b_v_*(*x, t*) = *b_e_*(*x, t*; *c*) now depends on *R_b_*(*t*). As a consequence, (29) is an integral equation to be solved for the unknown *R_b_*(*t*) concurrently with the solution of the IBVP (3)-(7) with 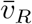 replaced by 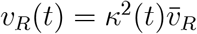.

### 3.1 Time Independent Steady State with Feedback

It has been shown in [24] that the extracellular model system without feedback has a unique steady state. We are interested here also in the corresponding steady state for the case with the spatially uniform non-local feedback on the signaling receptor synthesis rate 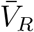 of the type characterized by (27). We denote the corresponding time independent steady state ectopic gradients by 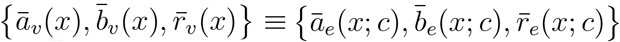 and the steady state robustness index 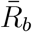 determined by

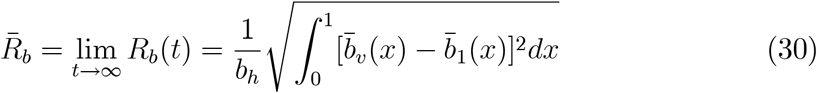

with *b_h_* appropriately taken to be *B*_1_(0), the LRO approximation for 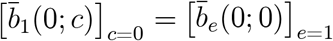. (Note that 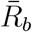 is positive unless *c* = 0 and *e* = 1.)

With *∂*()/*∂t* = 0, the governing equations and boundary conditions for our extracellular model with feedback can again be reduced to a BVP for *ā_v_* alone:

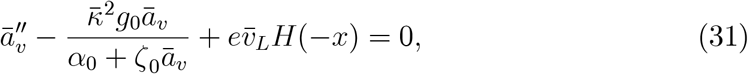

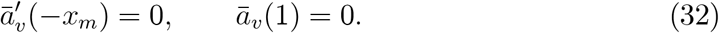

where *α*_0_ and *ζ*_0_ as previously given in (12) and

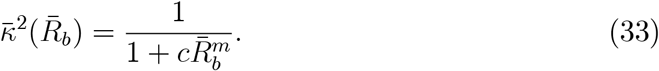

The corresponding signaling ligand and unoccupied receptor concentrations are given in terms of *ā_v_*

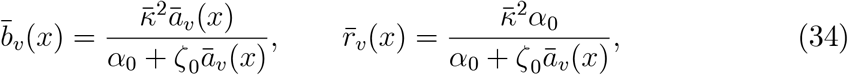

respectively. The solution for 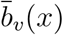 with 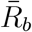 as an unknown parameter is then used in (30) for the determination of 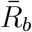.

Sample solutions of the BVP (31)-(32) and the signaling gradient 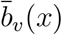 have been calculated and analyzed in [45]. Here, we complement the results there by establishing the existence, uniqueness and positiveness of the robustness index 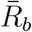 for the particular feedback problem. The method may be used to establish similar results for problems that include the effects of non-receptors in later sections.

To be concrete, we take *x_ℓ_* = 1, *x_h_* = 0, and *b_ℓ_* = 0 henceforth so that

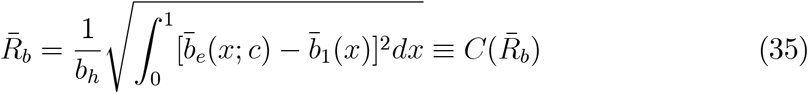

As indicated previously, we take *b_h_* to be given by (24). We now work with (35) to show first that 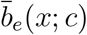 is a decreasing function of 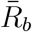 and then use the result in

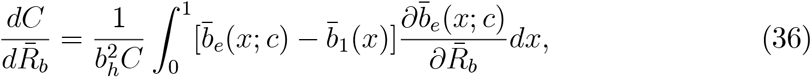

to establish the existence, uniqueness and positivity of 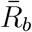.

Upon differentiating all relations in the BVP for *ā_v_*(*x*) = *ā_e_*(*x*; *c*) partially with respect to 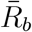, we obtain

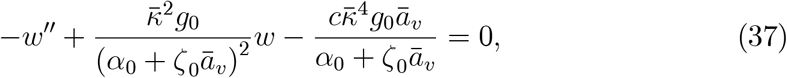

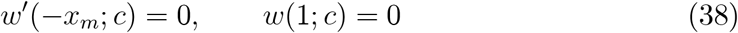

where

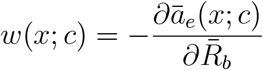

Clearly, *w_ℓ_*(*x*; *c*) = 0 is a lower solution of the BVP for *w*(*x*; *c*) given

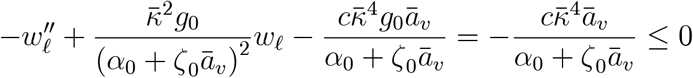

and (38). As an upper solution, we take

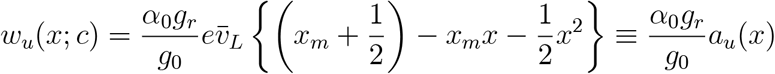

so that the boundary conditions for *w*(*x*) are satisfied and *w_u_*(*x*; *c*) ≥ 0 for all *x* in [−*x_m_*, 1]. Recall from [24] that *a_u_*(*x*) is an upper solution for *ā_v_*(*x*) so that *a_u_*(*x*) > *ā_v_*(*x*). With *ζ*_0_ = *O*(10^-1^) while 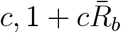 and 1 + *ζ*_0_*ā_v_*/*α*_0_ are *O*(1) quantities, we have

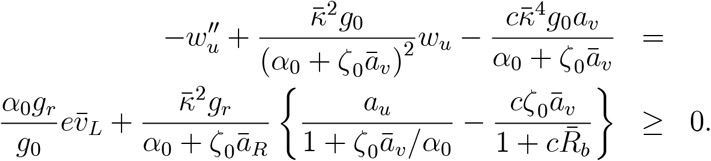

Then the monotone method in [38] implies that *w*(*x*; *c*) exists, is unique and nonnegative so that

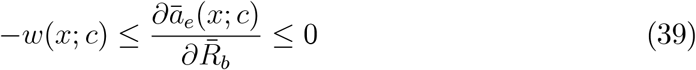

This development above leads to the following proposition on the non-positivity of the marginal value 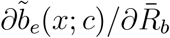.

#### Lemma 4

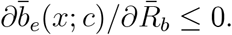

**Proof.** Upon differentiating the expression for 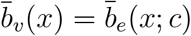 in (34) partially with respect to 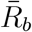, we obtain

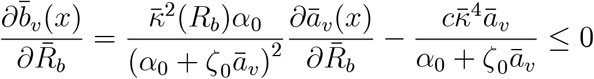

in view of the second inequality of (39).

#### Proposition 5

*A positive solution of* 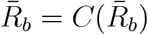 *exists and is unique*.

**Proof.** Together with 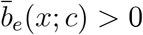, Lemma 4 implies 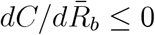 as long as 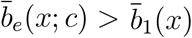 (see (36)). It follows that a solution exists for 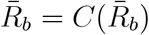. Itisuniquesince 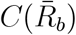 is monotone decreasing. It is positive since 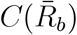 is nonnegative so that 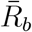 is bounded below with 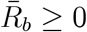.

### 3.2 Low Receptor Occupancy

The LOR state approximation for morphogen systems has been found useful for an understanding of the effects of various feedback mechanisms. The steady state approximation for the present feedback process has been obtained in [45]. The results are summarized below to be referenced later in the study of the effects of multi-feedback system involving non-receptors on robustness. In a state of *low receptor occupancy* prior to and after ligand synthesis enhancement so that

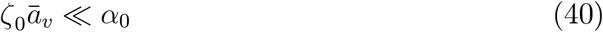

(including the special case where *c* = 0 and *e* = 1 so that *ā_v_*(*x*) reduces to *ā*_1_(*x*) = [*ā_e_*(*x*, 0)]_*e*=1_), it was shown in [45] that *ā_v_*(*x*) may be approximated by *eA_v_*(*x*) with

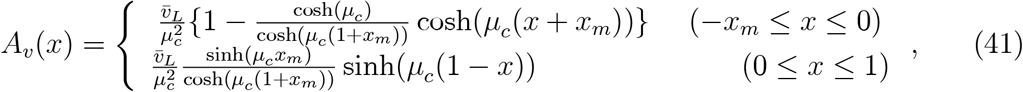

with

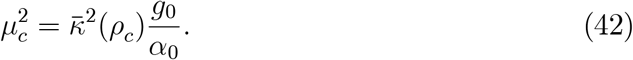

where *ρ_c_* is the LRO approximation of 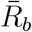 calculated from (30) using *eA_v_*(*x*) and *A*_1_(*x*, 0) for *ā_v_*(*x*) and *ā*_1_(*x*) = [*ā_e_*(*x*, 0)]_*e*=1_, respectively. The LOR solution *eA_v_*(*x*) is expected to be an accurate approximation of the exact solution *ā_v_*(*x*) and reduces to the LRO wild-type ligand concentration *A*_1_(*x*) when *c* = 0 and *e* = 1.

The corresponding LOR signaling gradient and free receptor concentration are given by

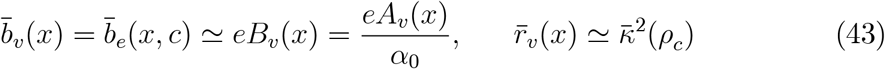

with

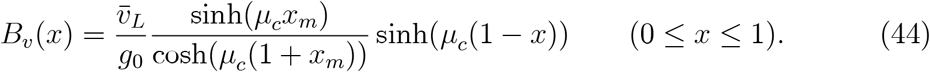

It should be noted that the dissociation rate constant is usually much smaller than the degradation rate constant with *f*_0_/*g*_0_ = *O*(10^-2^). To simplify our discussion, we have adopted the simplifying approximation *α*_0_ = (*f*_0_ + *g*_0_)/*h*_0_ ≃ *g*_0_/*h*_0_ so that

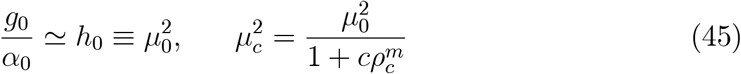

as in [45].

For *e* > 1, the yet unknown LOR robustness index *ρ_c_* is to be determined by the LRO approximation of (30)

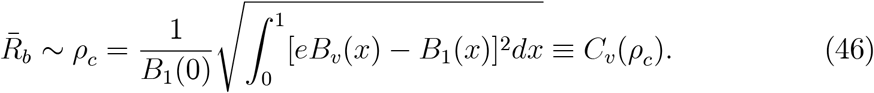

Given (19), (20) and (44), the right hand side depends on *ρ_c_* through *B_v_*(*x*); hence (46) is an equation for 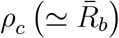:

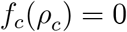

where

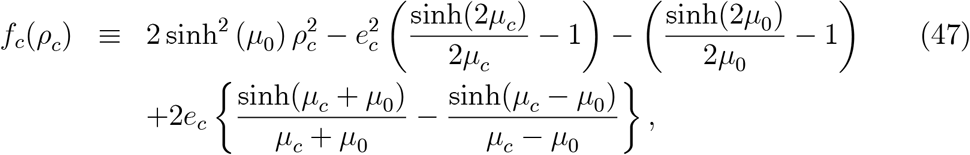

with

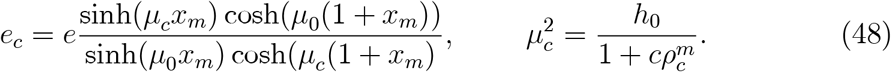

Even without an explicit solution for *f_c_*(*ρ_c_*) = 0, we see from (46) that *ρ_c_* is necessarily positive when *e* > 1 and therewith 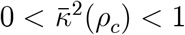. It follows from

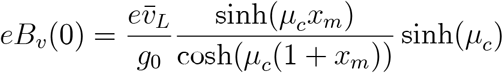

that the order of magnitude of *B_e_*(0, *c*) = *eB_v_*(0) is not changed in any significant way by the presence of the adopted feedback. With 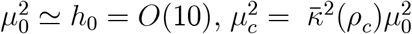 and 0 < *x_m_* ≪ 1, we can work with the approximate expression estimate

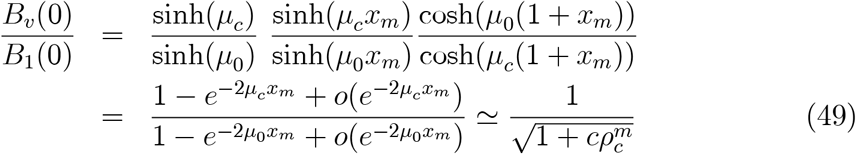

to get

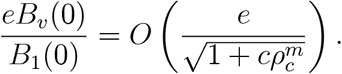

with *ρ_c_* < 0.2 for a robust gradient.

In addition, the negative feedback (27) also leads to a less convex modified ectopic signaling gradient since *μ_c_* < *μ*_0_ whenever *ρ_c_* > 0. We summarize the development above by the following observation:

#### Conclusion 6

*When both the wild-type and ectopic gradients are in a state of LRO, the negative feedback mechanism (27) on receptor synthesis rate is not particularly effective for promoting robust development*.

### 3.3 Numerical Solution for the General Case

In order to confirm the usefulness of the LOR solution for the problem, we have reconfigure the integro-differential equation problem for *ā_v_* as a BVP for a system of ODE solved it by available BVP solver. Numerical results are reported below for a system characterized by the parameter values shown in Table I.

The biological implication of the resulting robustness index is not particularly gratifying. The values of *ρ_c_* and 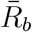 shown in the Table differ by less than 1% for all five values of *c*. Their values for *c* =1 are well above the acceptable level of 0.2, a rather modest requirement set arbitrarily in [26]) for robustness. This is hardly surprising given the estimate for the amplitude of the explicit solution *eB_v_*(0) for the LRO approximation of 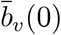 in (49) being independent of feedback. Comparing the accurate numerical solution for 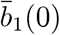 and 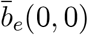 with 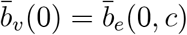 shown in Table I shows that the latter is very much closer to 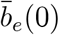 than 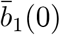. More specifically, 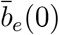 and 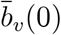 are roughly double the magnitude of 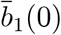, confirming the ineffectiveness of the feedback 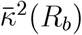 for *c* =1.

The LOR solution *eB_v_*(*x*) also shows that increasing the value of the parameter c to larger than 1 would further distort the shape of the gradient and ameliorate the amplitude reduction of the ectopic gradient through the terms involving hyperbolic functions with increasingly smaller *μ_c_*. Accurate numerical solutions for the general case (not in a state of LRO) in Table I are qualitatively similar to the LRO results with the robustness index decreasing rather gradually as c increases. Figure 1 shows graphs of 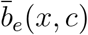 for *c* = 0, 1, 2 and 4 for comparison with 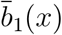 to further illustrate these observations. Note that for all the figures, the labels are generically 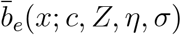 with *Z* = *η* = *σ* = 0 for Figure 1 (since these parameters do not appear until later sections) and 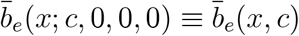.

**Figure 1:** Spatially uniform negative feedback on receptor synthesis rate.

It is necessary then to look elsewhere for a more effective feedback instrument to attain robust development with respect to an ectopic Dpp synthesis rate. There are a number of different such instruments available for this purpose. In the next few sections, we will focus on only one of these, namely, the role of non-receptors on robust development to support the findings for model 3 in [26]. This intermediate step also serves to pave the way to an appropriate multi-feedback mechanism for promoting robust signaling gradients.

As we routinely work with normalized quantities, we give below for subsequent references the normalized parameters corresponding to the actual biological parameter values listed under Table I above to be used for computing various solutions:

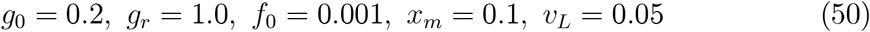

Note that the normalized value of receptor synthesis rate does not appear explicitly or used in various calculations. It is involved in various normalized quantities such as *v_L_* and will not be given here.

## 4 Effects of Non-receptors

The existence of inhibiting non-receptors and the associated feedback process are well documented for the BMP family ligands that includes the Dpp ligand of interest here (see [47, 34] and elsewhere). Known non-receptor type inhibitors include noggin, chordin, dally, follistatin, sog *and* various heparan sulfate proteoglycans. They are ubiquitous during wing imaginal disc and other biological developments (see [3, 5, 17, 20, 31, 37, 46, 48, 50] and references cited earlier). For the purpose of establishing robust signaling gradients, we are interested in the effects of non-receptors as an inhibiting agent on such gradients. The impact of non-receptors on the (time independent) steady state of a signaling gradient has been investigated theoretically by analysis and numerical simulations in [26, 29, 30, 44, 27] for models with a temporally and spatially uniform non-receptor synthesis rate ab initio. To the extent that inhibitors for promoting robust gradients are expected to be induced by ectopicity (as it would be a part of the wild-type development otherwise), the introduction of non-receptors in these models may be interpreted as a non-receptor synthesis rate of the form

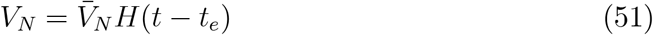

where *t_e_* > 0 is the instant of the onset of genetic or epigenetic perturbations. In this section, we examine the consequences of non-receptors generated in this way principally to delineate similar results obtained in [26] for (51) in the presence of a negative feedback on receptor synthesis rate. The usefulness of non-receptors for promoting signaling gradient robustness will be investigated for robustness index induced feedback on non-receptor synthesis in the next section.

To examine the results in [26], we take as a reference level of non-receptor concentration 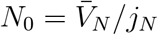 with

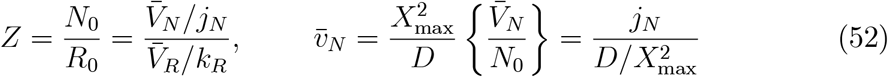

where *j_N_* is the degradation rate constant for the unoccupied non-receptors. Similar to signaling receptors, (normalized) free non-receptor concentration *n*(*x, t*) is also bound reversibly to Dpp ligand (to form normalized bound non-receptors of concentration *c*(*x, t*)),

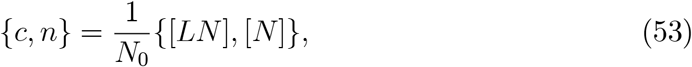

with a normalized binding rate constant *h*_1_*a* (for binding between Dpp and non-receptors of concentration [*N*]), non-receptor-mediated degradation rate constant *g*_1_ (for degradation of Dpp-non-receptor complexes [*LN*]), dissociation rate constant *f*_1_ (for dissociation rate of Dpp-non-receptor complexes) and free non-receptor degradation rate constant *g_n_* (for degradation of unoccupied non-receptors):

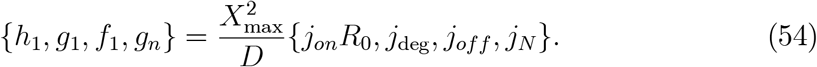

with *f*_1_ ≪ *g*_1_ so that *α*_1_ ≃ *g*_1_/*h*_1_.

In terms of these normalized quantities, we have the following IBVP for the five normalized unknowns *a, b, r, c* and *n* [44]:

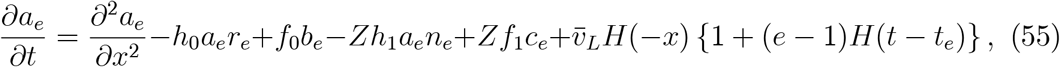

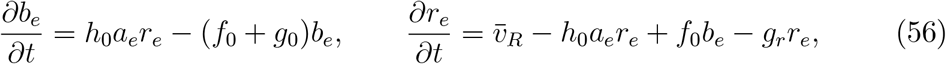

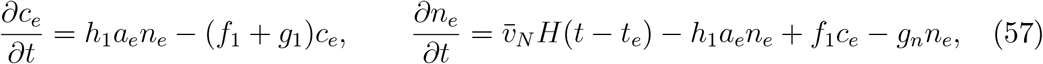

with the end conditions

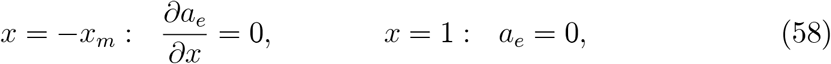

for *t* < 0, and (for *t_e_* < 0) the initial conditions

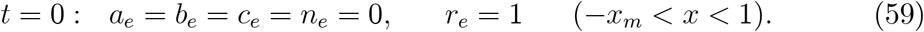

As before, the parameter *e* ≥ 1 is a measure of the ectopicity of the enhanced Dpp synthesis rate induced by a sustained genetic or epigenetic perturbation initiated at *t_e_* > 0. In this section, our first goal is to investigate whether the presence of a sufficiently high non-receptor synthesis rate may make the eventual signaling morphogen gradient [*LR*] insensitive to an ectopic Dpp synthesis rate. Since we are interested in the steady state behavior for large *t*, the initial conditions (59) will not have a substantive role in our analysis.

### 4.1 Time-Independent Synthesis Rates

When all synthesis rates for ligands, receptors and non-receptors are time-invariant, the solution of the new five-component model is expected to tend to a steady state denoted by 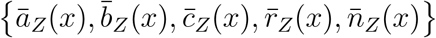 (alternative notations for 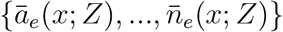 for brevity when appropriate). For this steady state solution, we have *∂*()/*∂t* = 0 so that the last four equations in (56) and (57) can be solved for 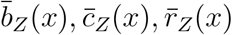 and 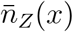 in terms of *ā_Z_*(*x*) to get (11) and

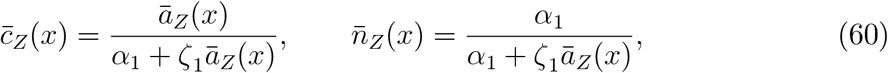

with

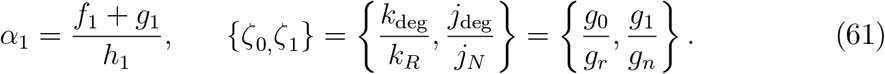

where *k*_deg_ and *j*_deg_ are the receptor- and non-receptor-mediated degradation rate constant, respectively, of bound Dpp complexes. The results are then used to obtain from the steady state version of (55),

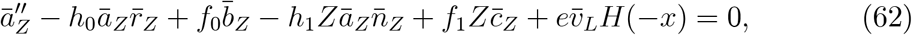

and (58) a BVP for *â_Z_*(*x*) = *ā_e_*(*x*; *Z*) alone:

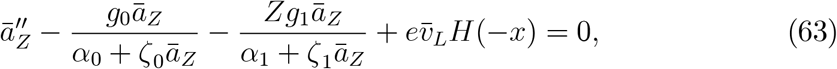

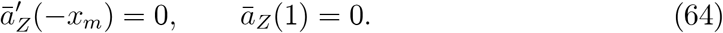

Evidently, *ā_Z_*(*x*) varies with *Z* (and occasionally written as *ā_e_*(*x*; *Z*)) which reduces to *ā_e_*(*x*) of the basic model for *Z* = 0, i.e., *ā_e_*(*x*) = [*ā_Z_*(*x*)]_*Z*=0_ = *ā_e_*(*x*; 0). Even if *e* = 1, *ā*_1_(*x*; *Z*) is different from the solution *ā*_1_(*x*) = *ā*_1_(*x*; 0), the free ligand concentration in wild-type development without non-receptors (corresponding to *t_e_* = ∞).

### 4.2 Low Receptor and Non-receptor Occupancy (LRNO)

For the signaling gradient to provide positional information that differentiates cell fates, the normalized concentration *b* = [*LR*]/*R*_0_ should not be nearly uniform (with a steep gradient adjacent to the absorbing edge or the ligand synthesis region). Positional indifference except for a steep gradient near the imaginal disc edge is not likely to occur if free ligand concentration is sufficiently low so that *α*_0_ + *ζ*_0_*ā_Z_* ≃ *α*_0_ and *α*_1_ + *ζ*_1_*ā_Z_* ≃ *α*_1_. In that case, the ODE (63) can be linearized with the corresponding approximate solution denoted by {*eA_Z_*(*x*), *eB_Z_*(*x*), *eC_Z_*(*x*), *eN_Z_*(*x*), *eR_Z_*(*x*)}. The relevant BVP has been reduced to a single ODE for *ā_Z_*(*x*) ≃ *eA_Z_*(*x*):

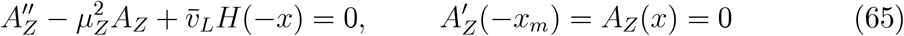

where

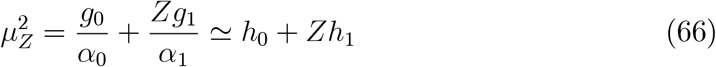

Its solution has been obtained in [26, 44] and elsewhere. With *f*_0_ ≪ *g*_0_ and *f*_1_ ≪ *g*_1_, we neglect (as in [45]) the effect of the dissociation rates and write *A_Z_*(*x*) as

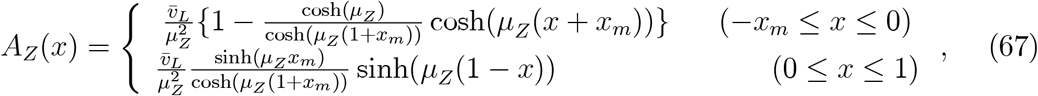

with 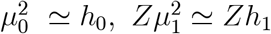 and

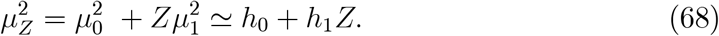

The corresponding signaling gradient is 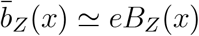 with

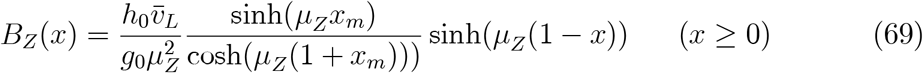

(keeping in mind *α*_0_ = (*g*_0_ + *f*_0_)/*h*_0_ ≃ *g*_0_/*h*_0_)and

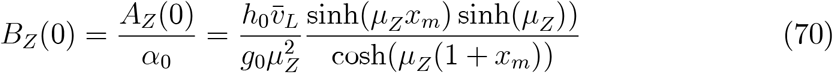

Note that *B_Z_*(*x*) depends on *Z* through *μ_Z_*.

A gradient system is said to be in a state of *low receptor and non-receptor occupancy* (LRNO) if

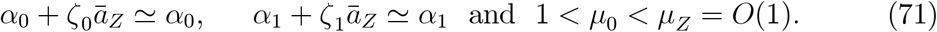

The gradient system is *biologically differentiating* if it is in a state of LRNO.

For *h*_1_ ≲ (*h*_0_)(> *g*_0_) so that 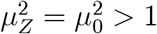, we have

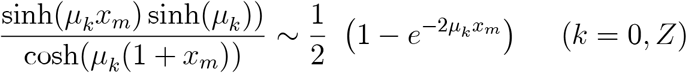

and

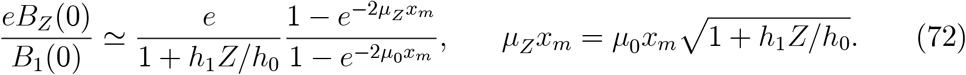

where 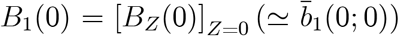. For *h*_1_ = *h*_0_ and *Z* = *O*(1), we have (with 0 < *x_m_* ≪ 1)

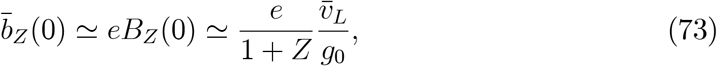

indicating that the amplitude of the ectopic signaling morphogen concentration (at *x* = 0) is reduced (approximately) to the wild-type concentration with *Z* = 1 for the empirically observable case of *e* = 2 (the amplification for Dpp synthesis rate associated with an increase of 5.9°*C* in ambient temperature). However, this does not mean that the ectopic gradient is reduced to the wild type since the shape parameter 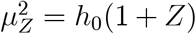 would be 2*h*_0_ so that the slope and convexity of the ectopic gradient would be changed substantially.

More generally, the ectopic concentration 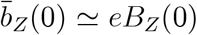 may evidently be kept to the wild-type level with a sufficiently high non-receptor synthesis rate so that 1 + *h*_1_*Z*/*h*_0_ ≥ *e*. However, for a given *h*_1_/*h*_0_ ratio, the required level of *Z* increases with *e* resulting in at least two effects that cause a distortion of signaling gradient shape and thereby work against the desired reduction of ectopicity of the signaling gradient. First, the ratio (1 − *e*^−2*μ_Z_x_m_*^)/(1 − *e*^−2*μ*_0_*x_m_*^) is significantly larger than 1 for larger *Z* and a larger *Z* is needed to reduce the amplitude of the gradient (than that for 1 + *h*_1_*Z*/*h*_0_ = *e*). Second, the ectopic gradient shape would be distorted even more severely by a larger *Z* since the gradient shape parameter 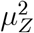 increases linearly with *Z*. The net effects from a particular non-receptor synthesis rate can only be seen from the corresponding value of the (LRNO approximation of the) robustness index, denoted by *ρ_Z_*. The dependent of *ρ_Z_* on *Z* will be calculated in the next subsection. Here, we settle for the following relatively conservative observation:

#### Conclusion 7

*For gradient systems in a steady state of low receptor and non-receptor occupancy with h*_1_/*h*_0_ = *O*(1), *the amplitude of their ectopic signaling gradient induced by the empirically observed synthesis rate amplification factor of e* = 2 *could be kept close to the wild-type amplitude (around x* = 0) *by a moderate non-receptor synthesis rate of Z* ≃ 1 *(as shown by the exact (numerical) solutions without the LRNO approximation in Figure 2)*.

**Figure 2:** Effects of Non-receptors

### 4.3 The LRNO Approximation of Robustness Index

As the presence of a fixed non-receptor concentration works both for and against robustness, whether or not the net effect is positive can only be seen from (the LRNO approximation *ρ_Z_* of) our adopted measure of robustness 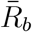. From (46), we have

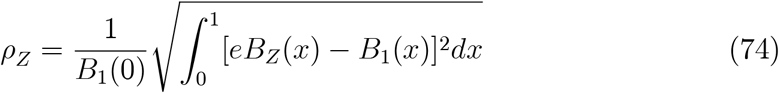

or

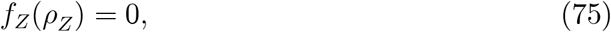

where *f_Z_*(·) is *f_c_*(·) as defined in (47) but with *μ_c_* and *e_c_* replaced by *μ_Z_* and

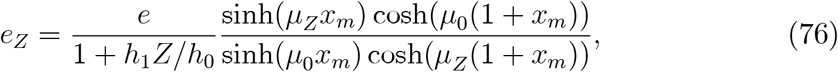

respectively.

Note that unlike its counterpart (46), the right hand side of (74) is independent of *ρ_Z_* so that it can be evaluated to get *ρ_Z_* (without having to solve a nonlinear equation as we had to do for *ρ_c_*). Upon writing (74) as

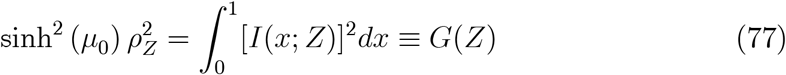

with

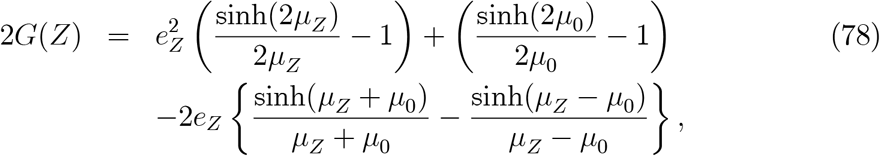

it is straightforward to plot *G*(*Z*) to see that 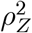 is a convex function of *Z* with a positive minimum at some minimum point *Z*_min_ > 0. Note that the convexity of *ρ_Z_* can also be seen analytically from the following properties of the integrand

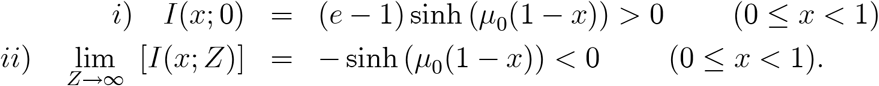

of (77) for any *x* in [0,1):

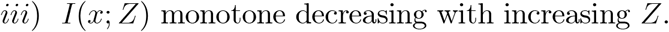

and, with 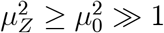,

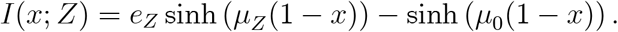

As a consequence, we have the following negative result similar to the corresponding finding in [26]:

#### Proposition 8

*The robustness of a signaling gradient in a steady state of LRNO deteriorates with increasing Z for Z* > *Z*_min_.

For the illustrating example of Table I, the optimal ratio *Z*_min_ that gives the smallest *ρ_Z_* is for *Z*_min_ ≅ 1.092 with *ρ*_min_ ≅ 0.0670 which is insignificantly below *ρ_Z_* = 0.0693… for *Z* = 1 but both are significantly below *ρ_Z_* = 0.1365… for *Z* = 2 (even if the latter is still below robustness threshold). Beyond an optimal level of *Z*, continuing increase in non-receptor synthesis rate would reduce the ectopic gradient concentration below the wild-type gradient 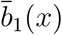 and worsen the corresponding shape difference, eventually to an unacceptable level of robustness.

Superficially, this suggests that there is no need to consider *Z* values higher than *Z*_min_ for a given problem, with *Z*_min_ ≃ 1 giving rise to an acceptable level of gradient shape distortion (as measured by our signal robustness index 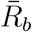). However, *Z*_min_ may have to be larger for another problem for which there is a need for a larger amplitude reduction factor 1 + *h*_1_*Z*/*h*_0_. For example, *Z* would need to be 3 or larger for the same problem with the larger ligand synthesis amplification factor (ectopicity) *e* = 4 (see (72) or (73)). This suggests that the induced non-receptor synthesis should be an increasing function *Z*(*e*) of the ectopicity factor *e*. A larger *Z* value would induce a much more severe (and probably unacceptable) signaling gradient shape distortion as seen from (69) and (68). These observations strongly suggest the following conclusion consistent with the simulation results of [26]:

#### Conclusion 9

*A non-receptor synthesis rate in the form of (51) and (55) does not offer a biologically realistic instrument for down-regulating ectopic signaling except for moderate ectopicity so that Z*_min_ ≃ 1. *In the latter case, robustness is sensitive to any further increase in non-receptor concentration*.

Accurate numerical results for the exact solution (without the LRNO approximation) 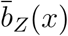 and 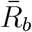 have been obtained for different (uniform) non-receptor synthesis rates (as characterized by the parameter *Z*) with additional parameters associated with the non-receptors assigned the following values in the illustrating example (also examined in [44]):

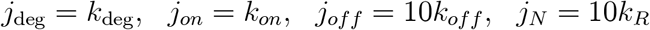

It is evident from the results reported in Table II that *ρ_Z_* and 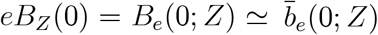 are quite accurate approximations of the corresponding numerical solutions for 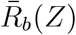 and 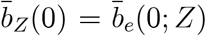 of the new model with (51). As such, the effects of non-receptors are pretty much delineated by the LRNO solution.

From either the computed LRNO solution or the numerical solutions for the original nonlinear BVP, we see that a moderate presence of non-receptors would generally bring 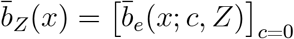 closer to 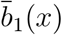 but generally renders 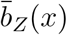 steeper and more convex than 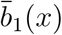 as shown in Figure 2. (Note that the notation 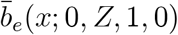 in Figure 2 corresponds to 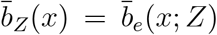.) The observations and conclusions above strongly suggest that the inhibiting mechanism (51) in not a realistic robustness promoting instrument. It is examined here not only to elucidate the findings by numerical simulations in [26] but also, with the help of the explicit analytical solution, to eliminate it as an instrument for promoting signaling gradient robustness. This tentative conclusion will be further strengthened by the developments in the next two subsections.

### 4.4 Addition of a Negative Feedback on Receptor Synthesis

The LRNO solution (69) shows that the distortion of the slope and convexity of ectopic signaling gradient by non-receptors is in the opposite direction relative to that induced by the feedback on signaling receptor synthesis rate given

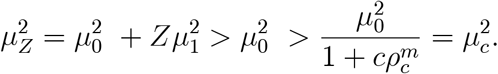

It would seem that we may be able to take advantage of this observation through a two-inhibitor model with receptor and non-receptor synthesis rates of the form (27) and (51), respectively, to more effectively promote robust gradients. The LRNO solution for such a model, denoted by *ā_e_*(*x*; *c, Z*) = *ā_RZ_*(*x*) ≃ *eA_RZ_*(*x*), is straightforward with

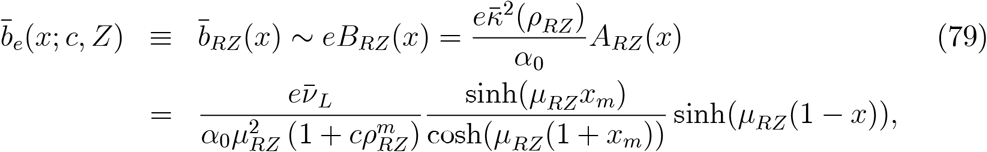

for 0 ≤ *x* ≤ 1, where *ρ_RZ_* is the LRNO approximation for the robustness index 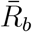 for the present model, and

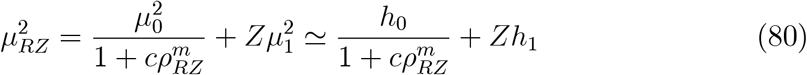

It is evident from the expression for 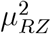 that its numerical value for a given gradient system is not substantially different from 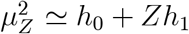 for *c* = *O*(1) since *ρ_RZ_* < *ρ_Z_* = *O*(10^-1^) for *Z* = *O*(1). This observation is confirmed by accurate numerical solutions for the new model (with and without the LRNO approximation) reported in Table III for the same sample problem as in Table I and II. The results show that there is not an appreciable reduction in the robustness index with *c* > 0 for *Z* = 1. The minimal effect of the signaling gradient itself is illustrated in Figure 3 for 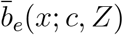. These observations are recorded as:

**Figure 3:** Effects of Negative Feedback on Receptor Synthesis for *Z* > 0

**Table III.** (*g*_0_ = *g*_1_ = 0.2; *h*_0_ = *h*_1_ = 10, *g_r_* = 1, *g_n_* = 10; *f*_0_ = 0.001, *f*_1_ = 0.01, *v_L_* = 0.05, *e* = 2)

| $Z$ | $c$ | $\bar{R}_b$ | $\rho_{RZ}$ | $\bar{b}_{RZ}(0)$ | $B_{RZ}(0)$ | $\bar{b}_1(0)$ |
| --- | --- | --- | --- | --- | --- | --- |
| 0 | 0 | 0.3943 | 0.3938 | 0.11517 | 0.1169 | 0.0579 |
| 0 | 1 | 0.3557 | 0.3565 | 0.10222 | 0.1062 | 0.0579 |
| 0 | 2 | 0.3269 | 0.3284 | 0.09407 | 0.0959 | 0.0579 |
| 1 | 0 | 0.0714 | 0.0693 | 0.07398 | 0.0739 | 0.0579 |
| 1 | 1 | 0.0619 | 0.0612 | 0.07550 | 0.0710 | 0.0579 |
| 1 | 2 | 0.0560 | 0.0566 | 0.06894 | 0.0688 | 0.0579 |
| 2 | 0 | 0.1298 | 0.1365 | 0.05597 | 0.0555 | 0.0579 |
| 2 | 1 | 0.1471 | 0.1547 | 0.05778 | 0.0497 | 0.0579 |
| 2 | 2 | 0.1683 | 0.1766 | 0.04463 | 0.0449 | 0.0579 |

#### Conclusion 10

*In the presence of a non-receptor inhibiting instrument of the form (51) and (55), the (steady state limit of a) spatially uniform non-local negative feedback (27) on receptor synthesis rate reduces the signaling gradient concentration and ameliorates the distortion of signaling gradient shape but not appreciably and only for Z* ≤ 1. *For Z* ≥ 2, *the robustness index* 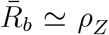 *actually deteriorates with the addition of the negative feedback on receptor synthesis rate*.

Similar to the case without the negative feedback on receptor synthesis, the results in Table III also show that too high a non-receptor-to-receptor synthesis rate ratio (*Z* ≥ 2 in our example) would cause an excessive reduction of signal morphogen concentration and too severe a shape distortion to result in an unacceptable robustness index. For *Z* ≥ 2 (needed for *e* > 2), the addition of feedback on receptor synthesis rate is actually deleterious to robustness for the illustrating example. The slight reduction of the shape distortion parameter 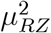 for a positive c, does not compensate for the considerably larger reduction of the amplitude reduction factor (that drives the amplitude of the signaling gradient at *x* = 0 to well below the concentration of the wild-type gradient at the same location). The corresponding graphs of 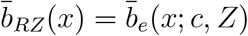 for different *Z* and *c* in Figure 3 clearly show why the robustness index 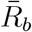 may eventually deteriorates with increasing *Z*.

To confirm, we have from

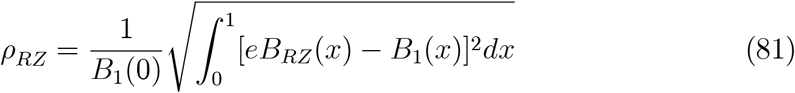

the equation for determining *ρ_RZ_*,

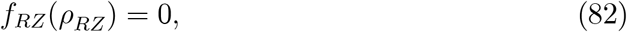

where *f_RZ_*(·) is *f_c_*(·) as defined in (47) but with *μ_c_* and *e_c_* replaced by *μ_RZ_* (with *m* taken to be 1 in (27)) and

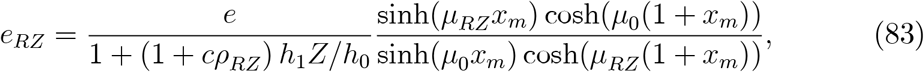

respectively. Solutions for some typical *ρ_Rz_* calculated from (82) are shown in Table III to confirm the observations made above and re-affirm Conclusion 9.

### 4.5 Root-Mean-Square Displacement Differential

With *μ_RZ_* and *e_RZ_* both depending on *ρ_RZ_*, the relation (82) is now a highly nonlinear equation for *ρ_RZ_*. Still, the expression for the new shape distortion factor *μ_RZ_* in (80) shows clearly that it is not possible for the negative feedback on receptor synthesis rate to reduce the shape distortion to an acceptable level if the non-receptor to receptor ratio *Z* should be much higher than 1. That this is not reflected in the computed values of *ρ_RZ_* in Table III suggests that the signaling robustness index *R_b_* is by itself not always an adequate measure of robustness. For this reason, we have also introduced in [26, 22] its companion robust index *R_x_* (denoted by *L* in [26]) that measures the root-mean-square displacement differential of the ectopic signaling gradient.

Let *x_e_*(*b*) and *x*_1_(*b*) be the location where the ectopic and wild-type gradients attains the value *b*, respectively, i.e., 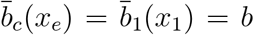. In steady state, the *displacement robustness index* 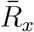 is defined by

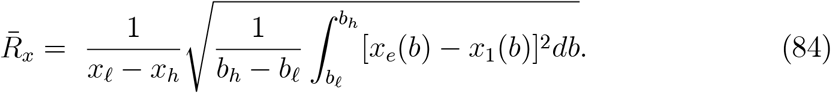

To minimize the effects of outliers, we may limit the range of *b* to be the interval (*b_ℓ_, b_h_*) with 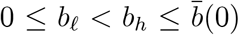. (We may take 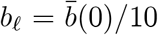 and 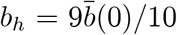 for instance.)

For gradients in a steady state of LRNO, the dependence of displacement Δ*x* = *x_e_*(*b*) − *x*_1_(*b*) on any feedback for a non-negative range of *x_e_* and *x*_1_ is through the expression

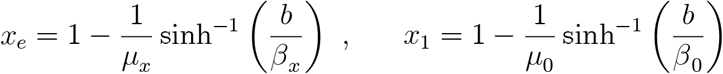

where 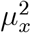 is 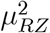 with *ρ_RZ_* replaced by *ρ_x_* (the LRNO approximation for 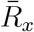) and

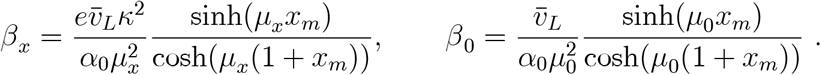

The LRNO approximation of 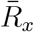 is then

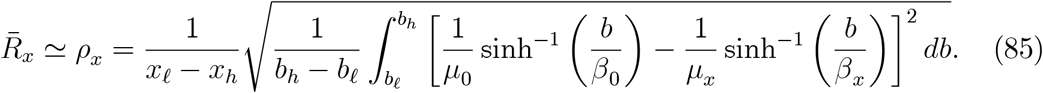

With 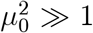, we may, to a good first approximation, work with the asymptotic values of these expressions to get

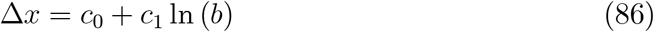

where

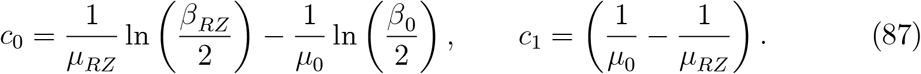

It follows that

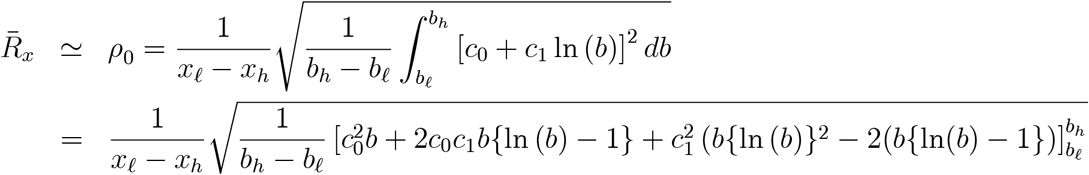

For sample calculations, we take *b_h_* = *b*(0) and *b_ℓ_* = *b*(1) ≈ 0 so that

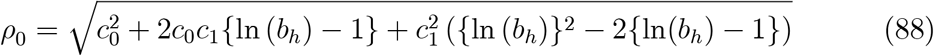

For more moderate values of *h*_0_ = *h*_1_ (with 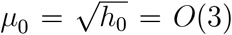), we would work with the exact inverse functions

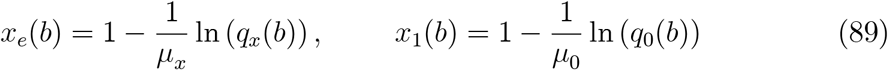

in (85) where

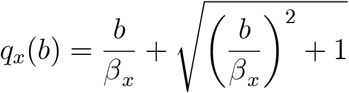

and *q*_0_(*b*) is *q_x_*(*b*) with *β_x_* replaced by *β*_0_. Table IV reports some typical results for the illustrating example of Tables I - III by the exact inverses (89):

**Table IV.** (*g*_0_ = *g*_1_ = 0.2; *h*_0_ = *h*_1_ = 10, *g_r_* = 1, *g_n_* = 10; *f*_0_ = 0.001, *f*_1_ = 0.01, *v_L_* = 0.05, *e* = 2)

|  |  |  |  |  |  |  |  |  |  |
| --- | --- | --- | --- | --- | --- | --- | --- | --- | --- |
| $(g_0 = g_1 = 0.2; \quad h_0 = h_1 = 10, \quad g_r = 1, \quad g_n = 10; \quad f_0 = 0.001, \quad f_1 = 0.01, \quad v_L = 0.05, \quad e = 2)$ | | | | | | | | | |
| $Z$ | 0 | 0 | 0 | 1 | 1 | 1 | 2 | 2 | 2 |
| $c$ | 0 | 1 | 2 | 0 | 1 | 2 | 0 | 1 | 2 |
| $\rho_{RZ}$ | 0.3938 | 0.3565 | 0.3284 | 0.0693 | 0.0612 | 0.0566 | 0.1365 | 0.1547 | 0.1766 |
| $\rho_x$ | 0.1987 | 0.2095 | 0.2159 | 0.0576 | 0.0575 | 0.0581 | 0.1454 | 0.1579 | 0.1737 |

With robustness measured by the new robustness index, the results for the illustrative example computed from the LRNO solution in Table IV now show clearly how *ρ_x_* deteriorates with increasing c even for typical values of *Z* around *Z*_min_ (while *ρ_RZ_* deteriorates with increasing c only for *Z* > *Z*_min_). They further strengthen Conclusion 9 and complement it with the following (instead of Conclusion 10):

#### Conclusion 11

*In the presence of the non-receptor inhibiting instrument of the form (51) and (55), the (steady state limit of a) negative spatially uniform non-local feedback (27) on receptor synthesis rate more readily exacerbates the ectopicity of the signaling gradient concentration when measured by the displacement robustness index ρ_x_*.

Given the different (and sometimes opposite) ways how the negative feedback on receptor synthesis rate impacts the two robustness indices of an ectopic gradient for *Z* ≥ 0, we should generally measure robustness by calculating both robustness indices before reaching any conclusion about the robustness of the signaling gradient.

## 5 Feedback on Non-receptor Synthesis Rates

### 5.1 The Model

Non-receptors as an inhibiting agent for down-regulating an ectopic gradient in the form (51) led to Conclusions 9 - 11 because the required synthesis rate has the consequence of increasing the shape distortion parameter severely for *Z* > 1 (which is needed for ectopicity factor *e* > 2). This would not be the case if the level of feedback depends on deviation from the wild-type gradient as measured by the robustness index. For this purpose, we consider a feedback instrument that also depends on the signaling robustness index of the form

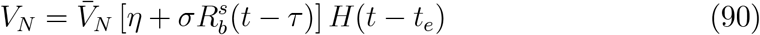

where the parameter *σ* and *s* characterize the strength and sensitivity of the feedback and the parameter *τ* pertains to a possible time delay on the effect of feedback (and *t_e_* > 0 is again the instant of onset of the genetic or epigenetic perturbations). The model in the previous section corresponds to *η* = 1 and *σ* = 0 while the case *η* = 0 and *σ* > 0 offers a positive feedback instrument for stimulating a non-receptor synthesis rate commensurate with the ectopicity measured by the robustness index *R_b_*. The subsequent development of the theory for this new feedback process is similar if the signaling robustness index *R_b_* is replaced by the corresponding displacement robustness index *R_x_* as defined in (84).

Anticipating a limiting time-independent steady state of signaling gradient, we expect

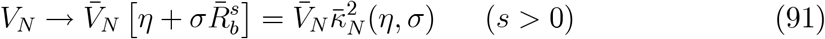

as the system approaches a time independent steady state. To maximize the effect of 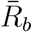 in the feedback mechanism (91), the parameter s will be set to 1 thereafter (similar to setting *m* = 1 in the feedback on receptor synthesis rate (27)). For that reason, the parameter *s* will not be displayed explicitly as a parameter of 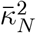 in (91) and elsewhere. Given Conclusion 9 and the fact that an instantaneously induced fixed synthesis rate is rather unrealistic, we limit our discussion for the *η* = 0 case. An ectopicity dependent *positive* feedback on non-receptor synthesis is also more consistent with experimental observations reported in references cited in the last section.

Upon modifying the model of Sec. 3 with the feedback on the receptor synthesis rate (27) to include the feedback (90), we reduce the new steady state problem to the following BVP for *ā_RN_*(*x*) = *ā_e_*(*x*; *c, Z, η, σ*):

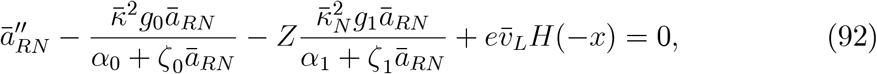

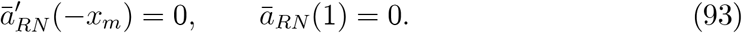

In a model with the feedback (91), the parameter *Z* will be set to 1 as the relative strength of non-receptor-to-receptor synthesis rate is readily set by the magnitude of *η* and/or *σ*.

Correspondingly, the various remaining gradients are given by

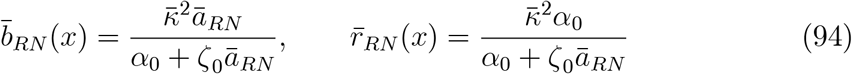

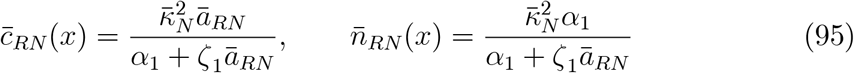

with 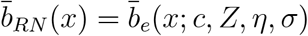, etc.

For a particular wing imaginal disc, it is clear from the model that the robustness of its development depends on the strength of the feedback on receptor synthesis rate characterized by the parameter *c* and the strength of the non-receptor synthesis rate characterized by the parameters *η* and *σ*. Similar to the numerical simulations of [26], exact numerical solutions of the nonlinear BVP (92)-(93) for our illustrating example have been obtained by typical BVP solvers to show typical effects of the three parameters *c, η* and *σ* (with *m* = *s* = 1) on the robustness index. These effects further confirm the benefits of setting *η* = 0 to be delineated more definitively by the explicit LRNO solution of the linearized problem in the next subsection.

### 5.2 Low Receptor and Non-receptor Occupancy

When both receptors and non-receptors are in a state of low occupancy, we may linearize the ODE (92) to get for *ā_RN_*(*x*) ≃ *eA_RN_*(*x*)

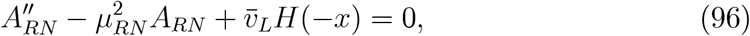

subject to the two end conditions (93) with

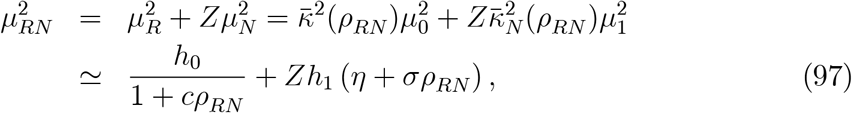

where *ρ_RN_* is the LRNO approximation of 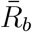 for the present model and we have made the highly accurate approximations 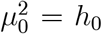 and 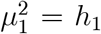 consistent with the simplification made in the earlier sections and in [45].

The exact solution for *A_RN_*(*x*) ≡ *A_e_*(*x*; *c, Z, η, σ*) is given by

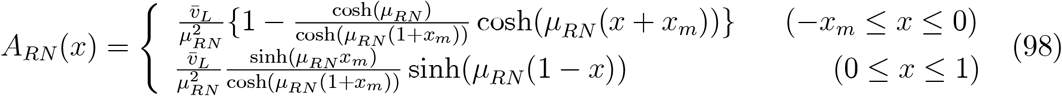

In the range 0 ≤ *x* ≤ 1, we have

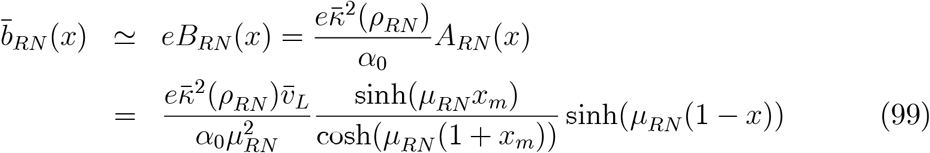

where 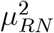 is as given in (97). From the expression

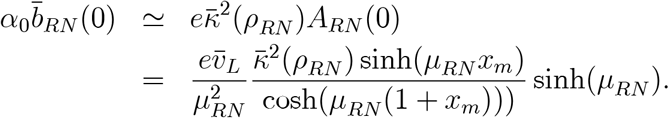

we obtain

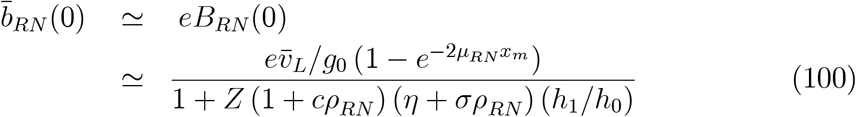

given *f_k_* ≪ *g_k_* for both *k* = 0 and 1. Correspondingly, we have

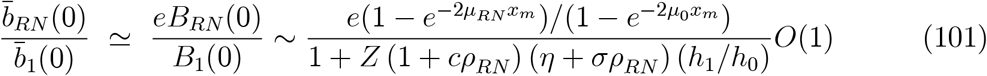

It is evident from (100)-(101) that, in the presence of non-receptors, a spatially uniform non-local positive feedback of the type (90) with *η* > 0 or *σ* > 0 (with *Z* = 1) reduces the signaling gradient concentration but also induces a shape change relative to that without any feedback. While this effect on the gradient shape counters that of the negative feedback on receptor synthesis, it may over-compensate the latter if the magnitude of *η* should be too large relative to what is needed to restore the signaling gradient to the corresponding wild-type gradient. With *η* = 0, the non-receptor concentration needed to achieve robustness can be adjusted by the level of ectopicity of the signaling gradient through the feedback strength parameter *σ*.

The net results of the different effects due to a positive feedback with *η* = 0 and *σ* > 0 is reflected in the LRNO robustness index

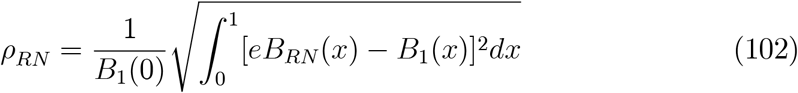

or

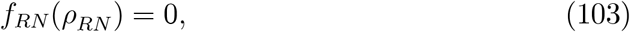

where *f_RN_*(·) is *f_c_*(·) as defined in (47) but with *μ_c_* and *e_c_* replaced by *μ_RN_* (with *m* and s both taken to be 1)and

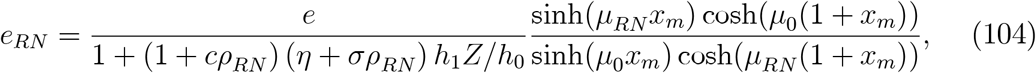

respectively. With *μ_RN_* and *e_RN_* both depending on *ρ_RN_*, the relation (103) is now a highly nonlinear equation for *ρ_RN_*.

### 5.3 Robustness Dependent Positive Feedback on Non-receptor Synthesis

Given Conclusion 9, we are interested here in a positive feedback on non-receptor synthesis that is robustness index dependent. The LRNO results for such a feedback alone is obtained from those of the previous section by setting *η* = 0 and *c* = 0. The signaling morphogen concentration 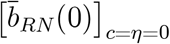, denoted by 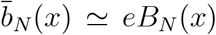, is given by

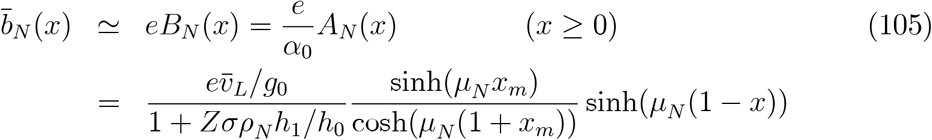

where *ρ_N_* = [*ρ_RN_*]_*c*=*η*=0_ is the LRNO approximation of the robustness index and

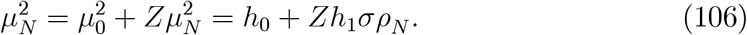

The signaling ligand concentration at *x* = 0 is

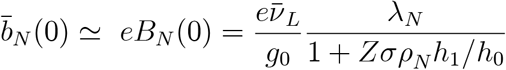

where

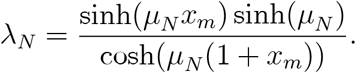

For the example used throughout this paper, we have *h*_1_/*h*_0_ = 1 and therewith

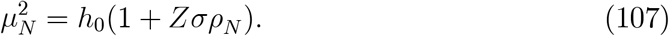

For *h*_0_ ≫ 1 (but *μ*_0_ = *O*(1) as in our illustrative example)

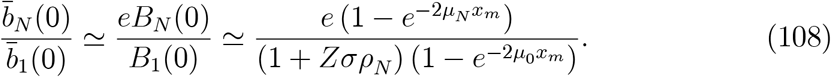

The LRNO approximation of the relative magnitude 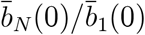 depends on the robustness index through the shape distortion parameter *μ_N_* and the amplitude reduction factor 1 + *Zσρ_N_h*_1_/*h*_0_. The LRNO approximation *ρ_N_* of the (signaling) robustness index is determined by

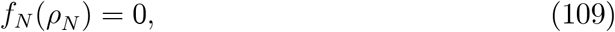

where *f_N_*(·) is *f_c_*(·) as defined in (47) but with *μ_c_* and *e_c_* replaced by *μ_N_* (with *m* and *s* both taken to be 1) and

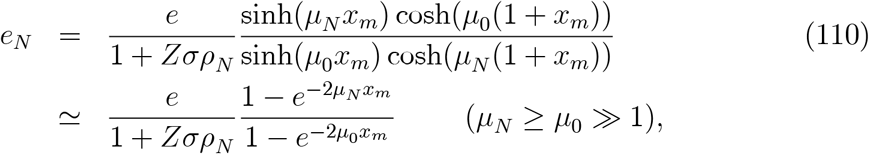

respectively (having specialized to the case *h*_1_ = *h*_0_).

The following observations are immediate from (110):

- The amplitude reduction factor 1 + *Zσρ_N_* is considerably smaller than the corresponding factor for the model with the non-receptor synthesis rate (51) when *Zσ* is *O*(*Z*) for the latter model (or *O*(*Zη*) in the feedback (91)) since *ρ_N_* is expected to be considerably less than unity.
- The shape distortion parameter *μ_N_* is considerably smaller than the corresponding parameter *μ_Z_* so that the shape of 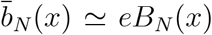 is less steep and less convex than 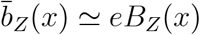.

By the first observation, the reduction of the signaling differential robustness index 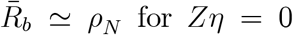 for *Zη* = 0 (and *Zσ* = 1) is expected to be considerably more modest than the corresponding reduction of 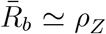 for *Zη* = 1 (and *Zσ* = 0). In particular, it barely meets the conservative threshold of *ρ_N_* ≤ 0.2 with *ρ_N_* = 0.1972 for *Zσ* = 2 for our illustrative example. This is understandable given the amplitude reduction factor is now (1 + *Zσρ_RN_*)^-1^ (for *η* = 0) instead of (1 + *Z*)^-1^ (for *η* = 1 and *σ* = 0) with *e*/(1 + *Z*) = 1 for *e* = 2 and *Z* = 1. We need *σρ_N_* = 0(1), i.e., *σ* = *O*(5), for the reduction to be comparable to the *η* =1 (and *σ* = 0) case. On the other hand, the shape distortion (for *η* = 0 and *σ* = 1) is now less severe with 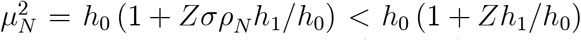 since *ρ_N_* < 1 for some degree of robustness. The actual benefit (or cost) for a *η* = 0 and *σ* = 1 feedback on nonreceptor synthesis rest on the net effect from the two opposite bulleted impacts above on the two robustness indices.

While this net effect may be case specific, it should be evident from the expression (107) that the impact of any increase in the synthesis rate ratio *Zσ* is much less than full (as it would be for the synthesis rate (51)) as only a fraction *ρ_N_* of *Zσ* is felt by the gradient system. Moreover, that fraction would be further reduced by the impact of the increase in *Zσ* (on reducing the robustness index). In other words, the impact of any increase in *Zσ* is doubly palliated (as long as the robustness index is less than unity), thereby reducing its effect on the shape distortion parameter *μ_N_*. In this sense, non-receptors as an inhibiting agent introduced through the feedback instrument (91) is stable, at least more so than through (51). This enables us to assert the following conclusion:

#### Conclusion 12

*The feedback instrument (91) with η* = 0 *is more stable and biologically realistic than η* > 0 *(and σ = 0) for down-regulating the ectopic signaling gradient*.

#### Remark 13

*For σ* > 0, *the addition of η* > 0 *would further decrease the amplitude reduction factor to* (1 + *Z*(η + *σρ*_RN_))^−1^. *However, the gain is offset by a more severe shape distortion resulting from* 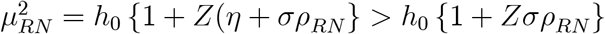.

### 5.4 An Effective Multi-Feedback Instrument for Robust Development

With *μ_Z_* > *μ_N_* ≥ *μ*_0_ > 1 ≥ *μ_c_*, the shape distortion induced by the feedback (91) with *η* = 0 is opposite to that by the negative feedback (33) on the receptor synthesis rate. By itself, the latter feedback has no impact on the amplitude reduction factor, administering concurrently, the two mitigating effects on shape distortion should help to lower the robustness index of the ectopic signaling gradient due to each feedback acting alone. To take advantage of this observation, we specialize the general results of Subsection 5.2 to the case *η* = 0 so that the positive feedback on non-receptor is robustness dependent. The signaling morphogen concentration 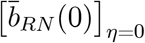,to be denoted by 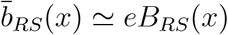, is given by

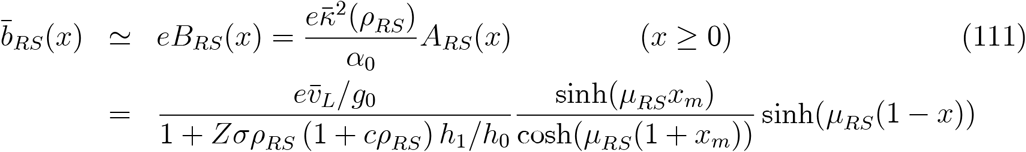

where *ρ_RS_* = [*ρ_RN_*]_*η*=0_ is the LRNO approximation of the robustness index and

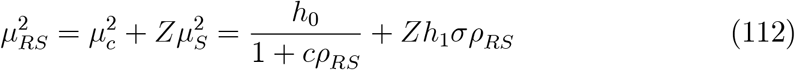

#### 5.4.1 The LRNO signaling morphogen concentration *eB_RS_* (0)

The LRNO signaling morphogen concentration at *x* = 0 is then given by

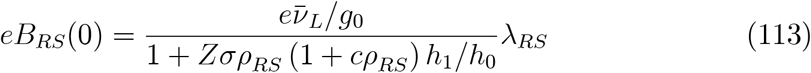

where

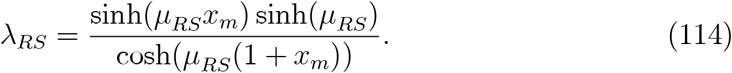

For the example used throughout this paper, we have *h*_1_/*h*_0_ = 1 and therewith

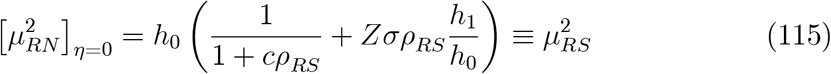

and

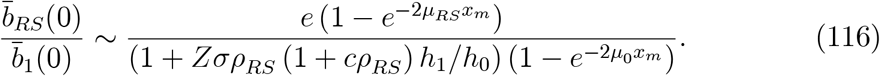

It is evident from (116) that the amplitude reduction factor is now larger than that for *c* = 0 so that it has the effect of reducing 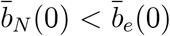. At the same time, we have 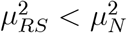 which reduces the shape distortion caused by *Zσ* > 0. Together, they enable us to conclude:

##### Conclusion 14

*Concurrent applications of the positive feedback on non-receptor synthesis (91) with η* = 0 *and a (moderate strength) negative feedback on receptor synthesis rate* 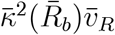 *promote more effectively robustness of an ectopic signaling gradient than either feedback acting alone*.

For a sufficiently large value of *c* > 0, 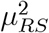 may be reduced to nearly 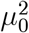 so that the resulting ectopic gradient shape would approach that of the wild type. With 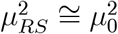 for *c* in the range that renders

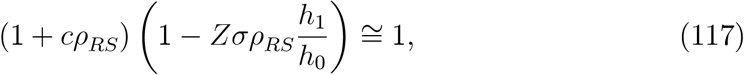

we have

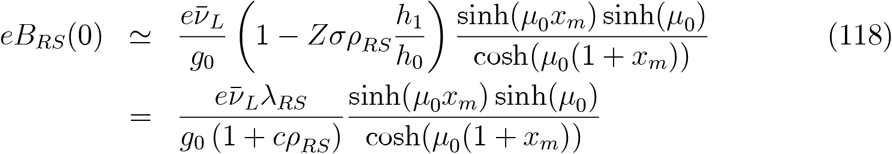

Whether the resulting ectopic signaling gradient is sufficiently close to the wild type can only be read from the corresponding two robustness indices.

#### 5.4.2 The LRNO Robustness Index

The LRNO approximation of the *differential signaling robustness index ρ_RS_* is determined by

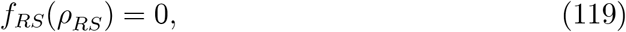

where *f_RS_*(·) is *f_c_*(·) as defined in (47) but with *μ_c_* and *e_c_* replaced by *μ_RS_* (with *m* and *s* both taken to be 1) and

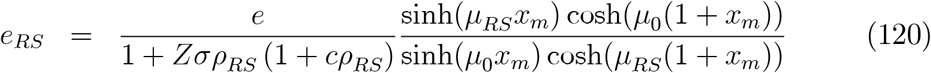

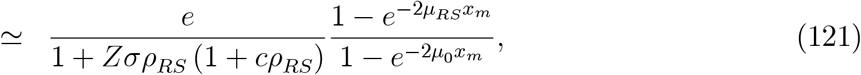

respectively (having specialized to the case *h*_1_ = *h*_0_). The relations (119) and (120) also apply to the corresponding *differential displacement robustness index ρ_x_* if we replace *ρ_RS_* in 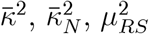, *e_RS_* and *f_RS_* by *ρ_x_* to be calculated from (85).

One effect from the addition of a negative feedback on signaling receptor synthesis rate on *ρ_RS_* is to increase the amplitude reduction factor to 1+*Zσρ_RS_* (1 + *cρ_RS_*) *h*_1_/*h*_0_ and thereby decreases the “amplitude” of *eB_RS_*(*x*) (see (113). Generally, this effect should reduce *ρ_RS_*. However, it is not difficult to see that

##### Lemma 15

*ρ_RS_ does not tend to* 0 *as c* → ∞.

**Proof.** Assuming the opposite so that *ρ_RS_* → 0 as *c* → ∞, there are three possibilities pertaining to the magnitude of *cρ_RS_*:

i. *cρ_RS_* → 0 with

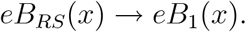

It follows that *ρ_RS_* → *ρ*_0_ > 0 as given in (26) contradicting the assertion to the contrary.

ii) 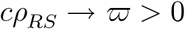 so that

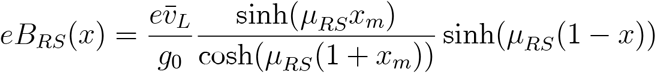

with

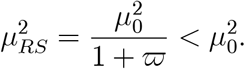

It follows that *eB_RS_*(*x*) is generally not *B*_1_(*x*) and *ρ_RS_* > 0 as *c* → ∞, contradicting the assertion to the contrary.

iii) *cρ_RS_* → ∞ but 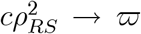, then we have 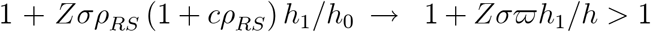 and 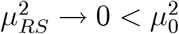 so that *ρ_RS_* must again be positive in the limit as *c* → ∞ and we again arrive at a contradiction.

As a consequence of the lemma above, we have the following observations on the robustness index:

- For a fixed value of *Zσ* (and *η* = 0), *ρ_RS_* may initially increase or decrease as c increases from 0 (for the same *Zσ* and *η* = 0) but must eventually reach a minimum and begin to increase with further increase in *c* tending to a finite limit *ρ*_∞_. Since the limit *ρ*_∞_ depends on the value of *Zσ*, the specific two-feedback mechanism is said to be *stable* (but not asymptotically stable).
- For any moderate value of *Zσ* (and *η* = 0) to result in a corresponding *ρ_W_*, the following inequalities on the shape distortion parameter and the amplitude reduction factor hold

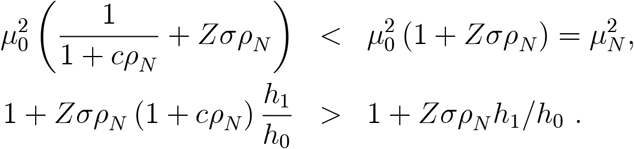 Hence, the addition of a moderate strength negative feedback on receptor synthesis rate should bring the ectopic signaling gradient closer to the wild type so that *ρ_RS_* < *ρ_N_*.
- For a fixed moderate value of *Zσ* so that *Zσρ_N_* < 1, a sufficiently large value of *c*, say *c*_1_,would render

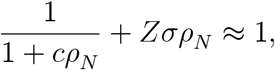

and keep the gradient shape close to the wild-type gradient shape. Increasing *c* well beyond *c*_1_ would reduce the shape parameter of the modified gradient well below unity, thereby would distort the gradient unduly in the opposite direction and work against robustness.

These results provide a specific realization of how an appropriate two-feedback combination of receptor and non-receptor synthesis rates may enhance both ectopic signaling ligand concentration reduction and shape-change amelioration:

##### Conclusion 16

*A multi-feedback instrument of the type (27) and (91) with η* = 0 *is both effective and stable for down-regulating the ectopic signaling gradient without distorting unacceptably the slope and convexity of the wild-type signaling gradient*.

The development above applies also to the displacement differential robustness index *ρ_x_*. As such the same qualitative conclusion may also be said about both indices. And both should be examined for robustness since they measure different features of the signaling gradient. Table V gives the two indices for the illustrative example for a range of *c* and *Zσ* showing the benefits of an appropriate combination of the multi-feedback instrument.

**Table V.** (*g*_0_ = *g*_1_ = 0.2; *h*_0_ = *h*_1_ = 10, *g_r_* = 1, *g_n_* = 10; *f*_0_ = 0.001, *f*_1_ = 0.01, *v_L_* = 0.05, *e* = 2, *η* = 0)

| $c \backslash Z\sigma$ | 0 | 1 | 2 | 4 | |
| --- | --- | --- | --- | --- | --- |
| 0 | 0.3938 | 0.2539 | 0.1972 | 0.1421 | $\rho_{RZ}$ |
| | 0.1987 | 0.1514 | 0.1249 | 0.0949 | $\rho_x$ |
| 1 | 0.3565 | 0.2263 | 0.1769 | 0.1289 | $\rho_{RZ}$ |
| | 0.2095 | 0.1526 | 0.1239 | 0.0930 | $\rho_x$ |
| 2 | 0.3284 | 0.2073 | 0.1626 | 0.1194 | $\rho_{RZ}$ |
| | 0.2159 | 0.1521 | 0.1223 | 0.0911 | $\rho_x$ |
| 4 | 0.2872 | 0.1815 | 0.1431 | 0.1059 | $\rho_{RZ}$ |
| | 0.2187 | 0.1483 | 0.1180 | 0.0872 | $\rho_x$ |

##### Remark 17

*Similar to the c* = 0 *case, the addition of η* > 0 *to the present two-feedback system (of c* > 0 *and σ* > 0*) would be a two edge sword: a further reduction of the amplitude reduction factor on the one hand, and an increase of the shape distortion factor* 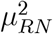 *the other hand. The former does not always promote robustness as it may down-regulate the signaling gradient to a level substantially below the wild-type gradient*.

**Figure 4:** Positive Feedback on Non-receptor Synthesis Only (*c* = *η* = 0)

**Figure 5:** Feedback on Receptor and Non-receptor Synthesis (*c* > 0, *η* = 0)

## 6 Concluding Remarks

Robustness with respect to an ectopic signaling gradient resulting from genetic or epigenetic perturbations requires one or more signaling activators to be down-regulated or inhibiting agents to be up-regulated by the enhanced signaling. This means the existence of some kind of feedback process in order to promote robustness. Feedback has long been seen as a mechanism for attaining robustness and specific feedback loops have been identified in the morphogen literature such as [10, 13, 14, 34, 36] and others). Though the conventional Hill function type negative feedback on receptor synthesis rate proves to be ineffective for this purpose [29, 19, 26], we have shown in [45] that appropriate multi-feedback instruments are capable of promoting robustness while individual feedback components of the multi-feedback arrangement fail to do so when acting alone.

Among the possible mechanisms for achieving robustness that appear biologically meaningful and realistic, non-receptors are known to be ubiquitous as an agent for down-regulating excessive signaling. That non-receptor would down-regulate signaling (and hence promote robustness) has already been established theoretically in [29, 30]. Yet numerical simulations in [26] feedback on non-receptors for promoting robustness often result in for gradient systems that are either still unacceptably ectopic or biologically unrealistic. Theoretical results of [45] suggests that a biologically realistic multi-feedback instrument involving feedback on non-receptors and one or more other known feedback processes exist and can be more effective in promoting signaling gradient robustness. The results of this paper confirm this anticipation and show an appropriate multi-feedback instrument involving non-receptors and receptor synthesis rate is in fact effective in promoting gradient robustness. They also suggest that other combinations of known feedback processes should be explored.

Understanding how robustness is attained by multi-feedback instruments is important not only to shed light on the reliability of developing gradient systems, but also to help explain the ubiquitous presence of elaborate regulatory schemes in morphogen systems.

## Supporting information

Supplemental_All-Figures

## 18 Acknowledgement

*The research of Q. Nie was partially supported by a NSF grant DMS1763272 and a grant from the Simons Foundation (594598, QN) awarded to UCI. The work of F. Y.M. Wan was supported NSF (UBM) DMS-1129008 awarded to UCI. The work of C. Sanchez-Tapia was supported by and NSF Grant awarded to Rutgers University*.

## Appendix

## A. Existence, Uniqueness, Non-negativity and Stability

### Theorem 19

*For positive values of the parameters g*_0_, *f*_0_, *h*_0_, 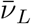 and 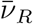, *there exists a unique, nonnegative solution ā_v_*(*x*) *of the BVP (31) and (32) linearly stable with respect to small perturbations. The corresponding concentrations* 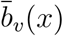 *and* 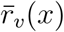 *can then be calculated from (34)*.

**Proof.** The existence proof is similar to that in [24] for the case without feedback. It suffices to produce an upper solution and a lower solution for the problem in order to apply the known monotone method in [38] (see also [2], [41]).

Evidently, *a_ℓ_*(*x*) = 0 is a *lower* solution since

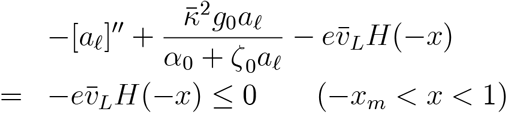

with

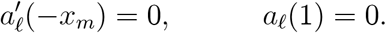

For an *upper* solution, consider

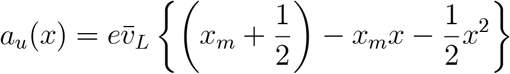

with 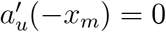 and *a_u_*(1) = 0. From (*i*) 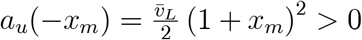, (*ii*) 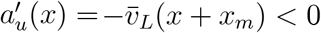 0 for *x* > –*x_m_*, and (*iii*) *a_u_*(1) = 0, we have

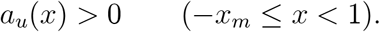

It follows from that 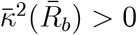 (see ()) that

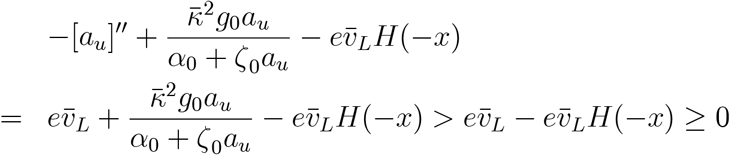

for –*x_m_* < *x* < 1 so that *a_u_*(*x*) is an upper solution for the BVP for *ā*(*x*). The monotone method assures us that there exists a solution *ã*(*x*) of the BVP (31) and (32) with

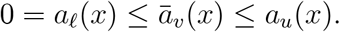

Since *a_u_*(*x*) is already known to be positive for – *x_m_* ≤ *x* < 1, *ā_v_*(*x*) must be nonnegative in the whole solution domain.

To prove uniqueness, let *a*^(1)^(*x*) and *a*^(2)^(*x*) be two (nonnegative) solutions and *a*(*x*) = *a*^(1)^(*x*) – *a*^(2)^(*x*). Then as a consequence of the differential equation (31) for *a*^(1)^(*x*) and *a*^(*2*)^(*x*), the difference *a*(*x*) satisfies the following differential equation:

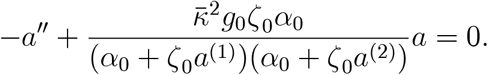

Form

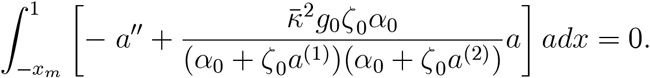

and integrate by parts. Upon observing continuity of *ā_v_*(*x*) and 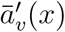, and application of the boundary conditions in (??), the relation above may be transformed into

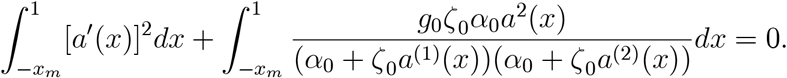

Both integrands are non-negative and not identically zero; therefore we must have *a*(*x*) ≡ 0 and uniqueness is proved.

Stability of the steady state solution with respect to small perturbations from the steady state can be proved by an argument similar to that used in [24] but will be omitted since it is not needed in subsequent developments.

## B. Monotonicity

As for the model analyzed in [24], free morphogen concentration *a_e_*(*x*) and the corresponding signaling morphogen gradient *b_e_*(*x*) can be shown to be (positive and) monotone decreasing in the open interval (–*x_m_*, 1). We show this by first ruling out the possibility of any extremum in that interval.

### Proposition 20

*Under the same hypotheses as those in Theorem 19, the nonnegative steady state concentration ā_v_*(*x*) *does not attain a maximum or minimum in* (0, 1) *and hence is monotone decreasing in that interval*.

**Proof.** First, it is easy to see that the nonnegative *ā_v_*(*x*) does not have an interior maximum in the interval 0 < *x* < 1. If it should have a local maximum at some interior point *x*_0_, then we must have (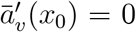 and) 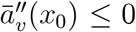. But since *ā_v_*(*x*) ≥ 0 and *v_L_*(*x*) = 0 in *x* > 0, we have

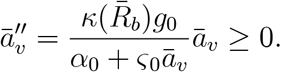

It follows that we must have 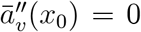 and therewith *ā_v_*(*x*_0_) = 0. Since *x*_0_ is a maximum point, we must have *ā_v_*(*x*) = 0 in 0 < *x* < 1. The continuity requirements imply 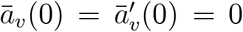. But it is impossible for any nontrivial solution of the ODE (63) to satisfy both of these conditions unless *ā_v_*(*x*) = 0 for all *x* in [–*x_m_*, 0] as well. But such a free morphogen concentration does not satisfy (63) in the interval (–*x_m_*, 0) where the normalized *Dpp* synthesis rate is a positive constant 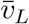. Hence *ā_v_*(*x*) does not have a maximum in (–*x_m_*, ∞).

Also, *ā_v_*(*x*) does not have a positive interior minimum. If it should have one at *x*_0_ (with *ā_v_* (*x*_0_) > 0), then it must have an interior maximum at some *x*_1_ > *x*_0_ in order for *ā_v_*(*x*) to decrease from *ā_v_*(*x*_1_) > 0 to *ā_v_*(1) = 0. But this contradicts the fact that *ā_v_*(*x*) does not have an interior maximum. There is still the possibility of a local interior minimum *ā_v_*(*x*_0_) = 0. With 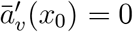 at the local minimum, we have *ā_v_*(*x*) ≡ 0 which does not satisfy the ODE (63) in the interval (−*x_m_*, 0).

Altogether, the solution *ā_v_*(*x*) of the BVP must be (nonnegative and) monotone decreasing from *ā_v_*(–*x_m_*) > 0 to *ā_v_*(1) = 0.

We can actually prove that the relevant morphogen concentrations are positive for *x* < 1 which we will need in subsequent development.

### Corollary 21

*Under the hypotheses of Theorem 19, the concentrations ā_v_*(*x*), 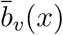 and 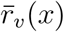 do not vanish in (–*x_m_*, 1).

**Proof.** Suppose *ā_v_*(*x*) vanishes at *x*_0_ in (–*x_m_*, 1) and hence attains a local minimum there (since *ā_v_*(*x*) is non-negative). But this contradicts Proposition 20 which asserts that *ā_v_*(*x*) does not have an interior minimum. That the remaining quantities do not vanish follows from (34) and (61).

University of California at Irvine

