## Supplemental_All-Figures for "Regulatory Feedbacks on Receptor and Non-receptor Synthesis for Robust Signaling"

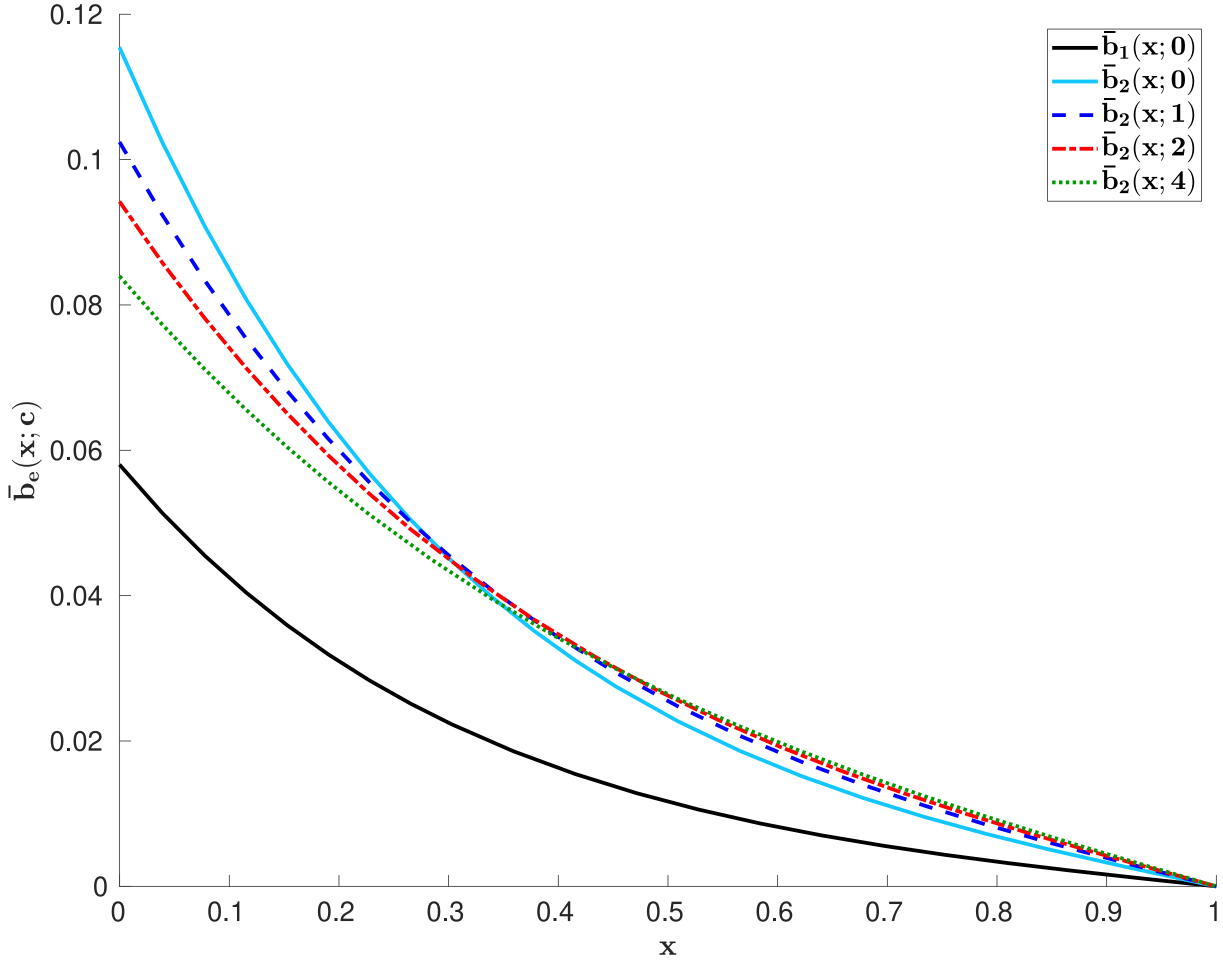


Figure 1: Spatially uniform negative feedback on receptor synthesis rate. Parameter values $g_1 = 0.2$, $h_1 = 10$, $f_1 = 0.01$, $g_n = 10$ and $v_L = 0.05$.

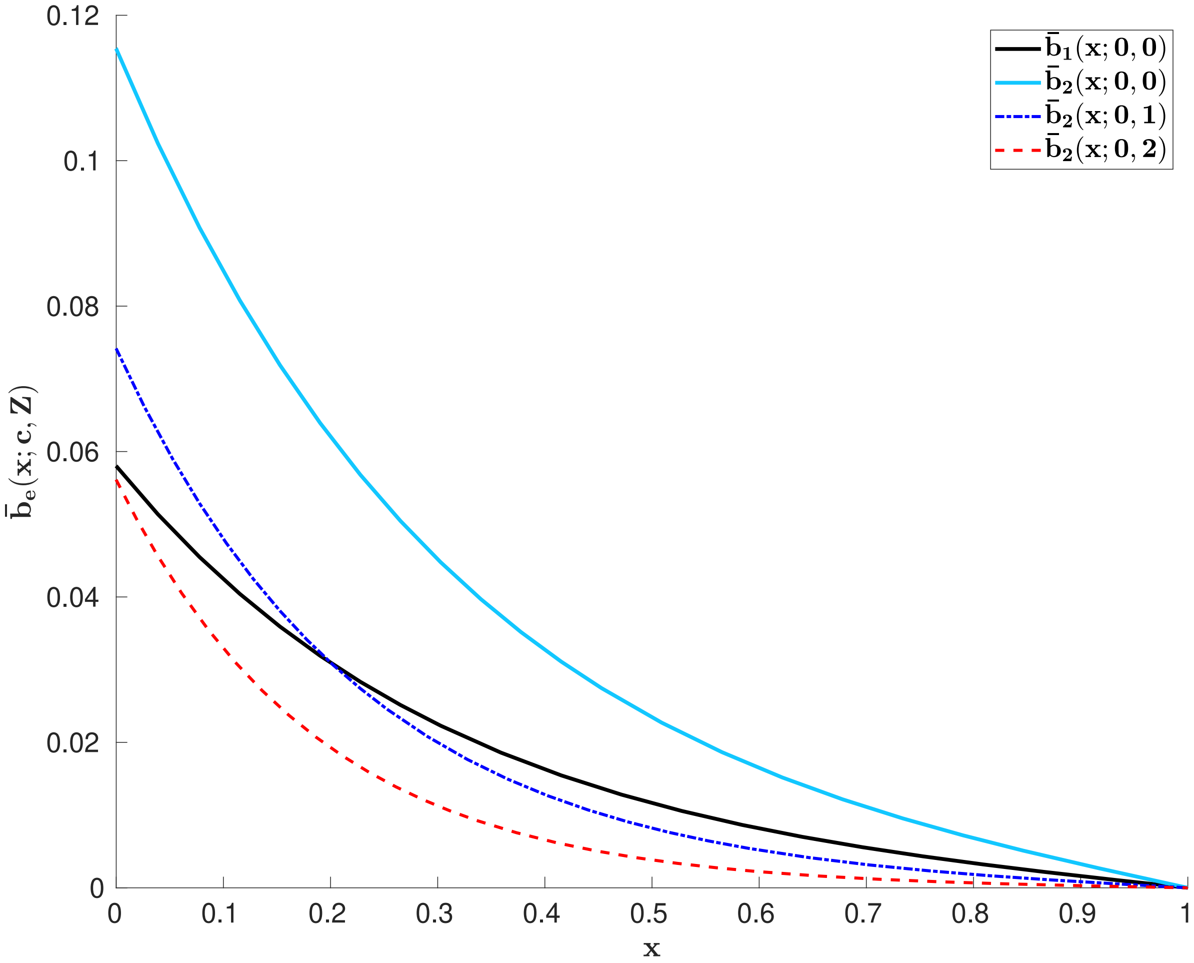


Figure 2: Effects of Non-receptors

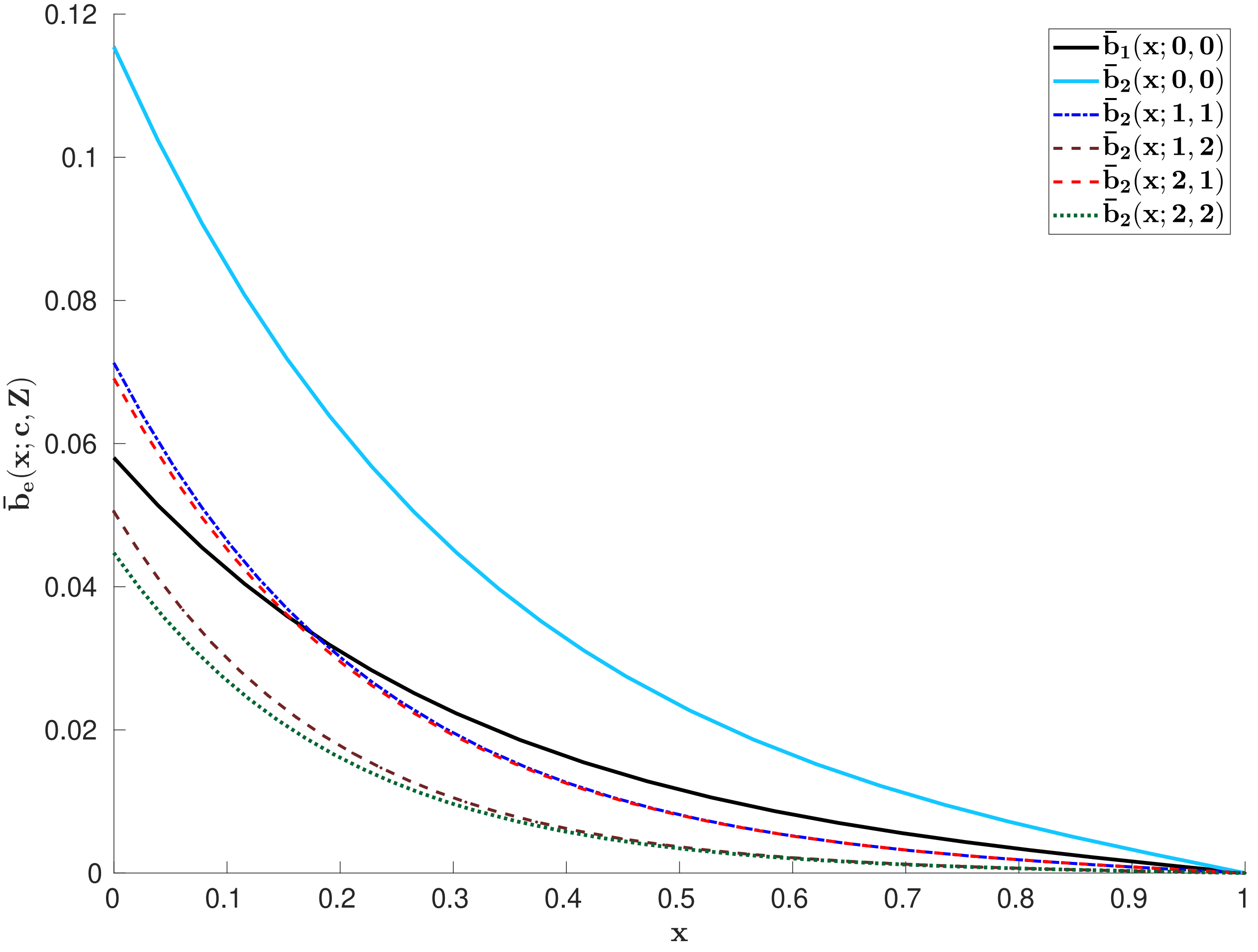
0.12
0.1
0.08
0.06
0.04
0.02
0
0.1
0.2
0.3
0.4
0.5
0.6
0.7
0.8
0.9
1
$\bar{\mathbf{b}}_{\mathbf{e}}(\mathbf{x};\mathbf{c},\mathbf{Z})$
x
$\bar{\mathbf{b}}_{\mathbf{1}}(\mathbf{x};\mathbf{0},\mathbf{0})$
$\bar{\mathbf{b}}_{\mathbf{2}}(\mathbf{x};\mathbf{0},\mathbf{0})$
$\bar{\mathbf{b}}_{\mathbf{2}}(\mathbf{x};\mathbf{1},\mathbf{1})$
$\bar{\mathbf{b}}_{\mathbf{2}}(\mathbf{x};\mathbf{1},\mathbf{2})$
$\bar{\mathbf{b}}_{\mathbf{2}}(\mathbf{x};\mathbf{2},\mathbf{1})$
$\bar{\mathbf{b}}_{\mathbf{2}}(\mathbf{x};\mathbf{2},\mathbf{2})$

Figure 3

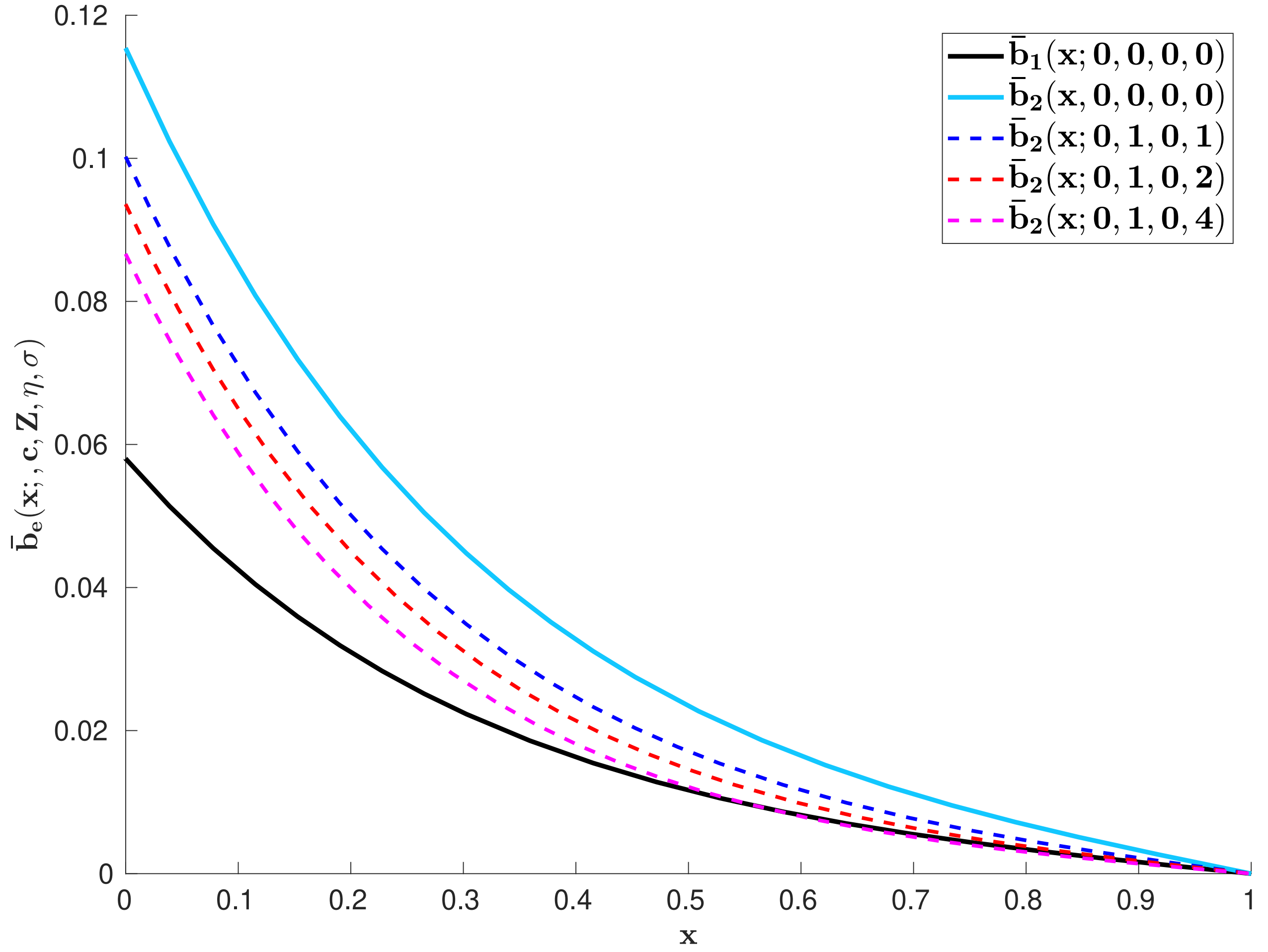
$\bar{b}_1(x; 0, 0, 0, 0)$
$\bar{b}_2(x, 0, 0, 0, 0)$
$\bar{b}_2(x; 0, 1, 0, 1)$
$\bar{b}_2(x; 0, 1, 0, 2)$
$\bar{b}_2(x; 0, 1, 0, 4)$
$\bar{b}_e(x; , c, Z, \eta, \sigma)$
x
0
0.02
0.04
0.06
0.08
0.1
0.12
0.1
0.2
0.3
0.4
0.5
0.6
0.7
0.8
0.9
1

Figure 4

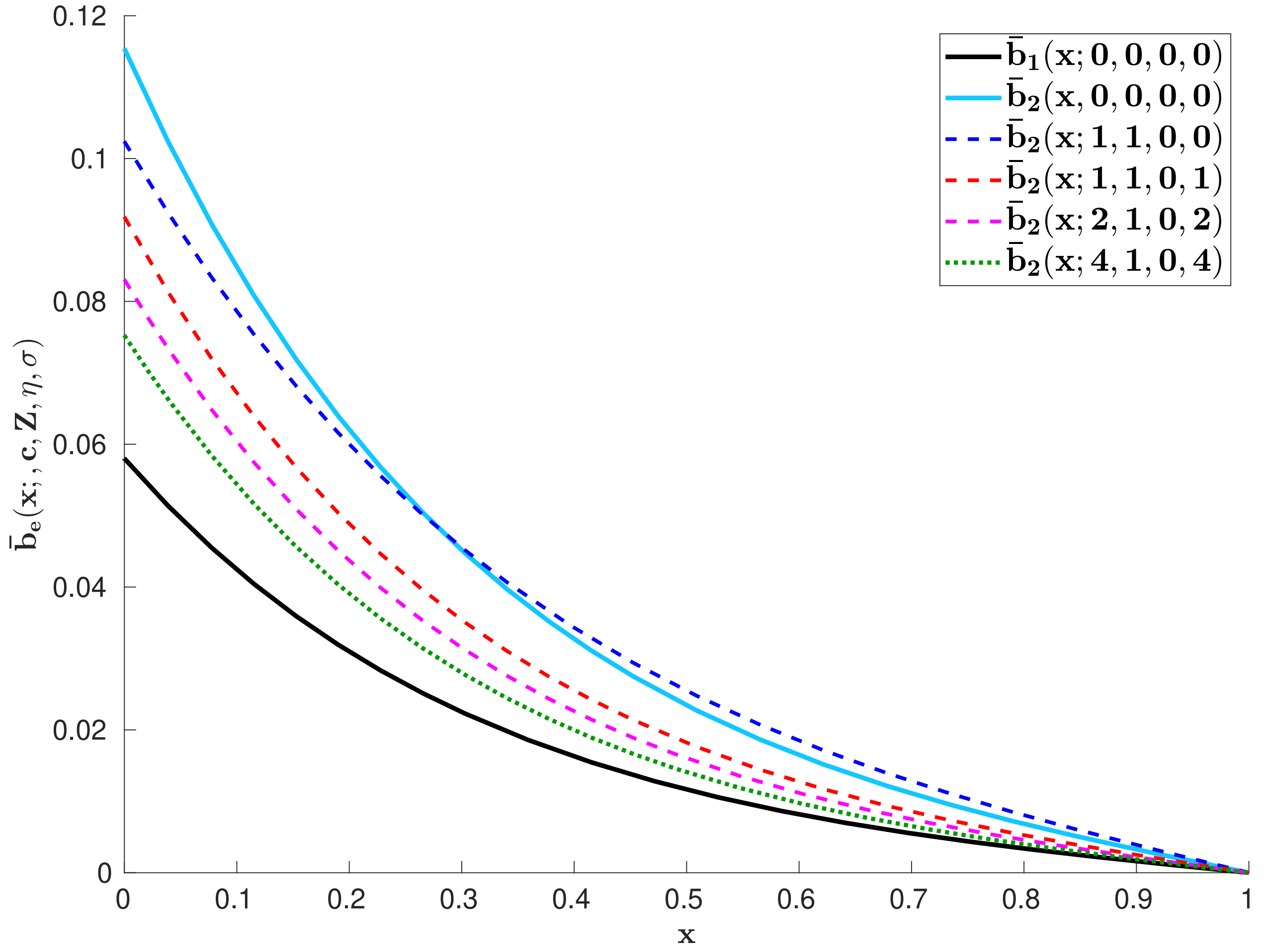
$\bar{b}_1(x;0,0,0,0)$
$\bar{b}_2(x,0,0,0,0)$
$\bar{b}_2(x;1,1,0,0)$
$\bar{b}_2(x;1,1,0,1)$
$\bar{b}_2(x;2,1,0,2)$
$\bar{b}_2(x;4,1,0,4)$
$\bar{b}_e(x;,c,Z,\eta,\sigma)$
x
0
0.02
0.04
0.06
0.08
0.1
0.12
0.1
0.2
0.3
0.4
0.5
0.6
0.7
0.8
0.9
1

Figure 5
